# A novel DNA binding mode, conformational change and intramolecular signaling in MutS upon mismatch recognition in replication DNA structures

**DOI:** 10.1101/2020.05.01.070730

**Authors:** Milagros Inés Ibáñez Busseti, Lucía Malvina Margara, Sofía Daiana Castell, Marisa Mariel Fernández, Emilio Luis Malchiodi, Guillermo Gabriel Montich, Virginia Miguel, Carlos Enrique Argaraña, Mariela Roxana Monti

**Affiliations:** Centro de Investigaciones en Química Biológica de Córdoba (CIQUIBIC), CONICET, Departamento de Química Biológica Ranwell Caputto, Facultad de Ciencias Químicas, Universidad Nacional de Córdoba, Ciudad Universitaria, Córdoba, Argentina; Structural and Computational Biology Unit, European Molecular Biology Laboratory, Heidelberg, Germany; Instituto de Estudios de la Inmunidad Humoral Profesor Ricardo A. Margni (IDEHU), CONICET, Departamento de Microbiología, Inmunología, Biotecnología y Genética, Facultad de Farmacia y Bioquímica, Universidad de Buenos Aires, Buenos Aires, Argentina; Instituto de Investigaciones Biológicas y Tecnológica (IIBIT), CONICET, Departamento de Química, Facultad de Ciencias Exactas, Físicas y Naturales, Universidad Nacional de Córdoba, Córdoba, Argentina

**Keywords:** MutS, mismatch repair, DNA replication, mutagenesis, Pseudomonas aeruginosa

## Abstract

MutS initiates mismatch repair by recognizing mismatches in newly replicated DNA. Specific interactions between MutS and mismatches within double-stranded DNA promote ADP-ATP exchange and a conformational change into a sliding clamp. Here, we demonstrated that MutS from *Pseudomonas aeruginosa* associates with primed DNA replication substrates. The predicted structure of this MutS-DNA complex revealed a new DNA binding site, in which Asn 279 and Arg 272 appeared to directly interact with the 3’-OH terminus of primed DNA. Mutation of these residues resulted in a noticeable defect in the interaction of MutS with replication DNA substrates. Remarkably, MutS interaction with a mismatch within primed DNA induced a compaction of the protein structure and impaired the formation of an ATP-bound sliding clamp. Our findings reveal a novel DNA binding mode, conformational change and intramolecular signaling for MutS recognition of mismatches within DNA replication structures.

## INTRODUCTION

Most living organisms possess the DNA mismatch repair (MMR) mechanism in order to avoid mutations and safeguard genomic stability. MMR corrects mismatches and insertion-deletion loops generated during DNA synthesis, enhancing the overall fidelity of DNA replication up to 1000-fold (Liu et al., 2017). MMR also plays critical roles in other cellular biological processes such as mitotic and meiotic recombination, and DNA-damage signaling. Defects in MMR factors are associated with a mutator phenotype, characterized by increased mutation and recombination rates, and result in an increased susceptibility to sporadic colorectal, endometrial and ovarian cancer in humans (Pecina-Slaus et al., 2020). Furthermore, germline mutations in MMR genes cause one of the most common forms of hereditary cancer, called Lynch syndrome, which is characterized by a predisposition to a spectrum of cancers, mainly colorectal and endometrial cancer (Pecina-Slaus et al., 2020).

The mechanism for repairing mutagenic replication errors has been deciphered using *in vitro* structural analysis and bulk and single-molecule biochemical experiments (Lenhart et al., 2016, Liu et al., 2017). The essential MMR steps comprise error recognition, discrimination of the nascent strand containing the error, strand excision and gap-filling DNA synthesis. These reactions are catalyzed by the core MMR machinery, including the highly conserved MutS and MutL proteins (and eukaryotic MutSα, MutSβ and MutLα), and accessory factors involved in multiple DNA metabolic pathways.

The mismatch sensing MutS protein, belonging to the ATP-binding cassette ATPase family, initiates MMR (Groothuizen and Sixma, 2016, Hingorani, 2016). ADP-bound MutS moves along the double-stranded DNA, searching for mismatches. Error recognition stalls the diffusion of MutS at the site of the mismatch and triggers the exchange of ADP by ATP (Acharya et al., 2003, Cho et al., 2012, Gorman et al., 2012, Gradia et al., 1999). ATP binding elicits the formation of a sliding clamp that dissociates from the mismatch in an ATP hydrolysis-independent manner and recruits MutL (Acharya et al., 2003, Gradia et al., 1999, Groothuizen et al., 2015, Liu et al., 2016). In enteric bacteria, including *Escherichia coli*, MutL activates the MutH endonuclease, which incises the unmethylated nascent strand in hemi-methylated d(GATC) sites (Acharya et al., 2003, Liu et al., 2016). The UvrD helicase unwinds the DNA starting from the nick, followed by strand degradation by single-stranded DNA exonucleases (Burdett et al., 2001, Dao and Modrich, 1998, Liu et al., 2019). The mechanism of strand-specific incision and excision remains poorly understood in bacteria and eukaryotes lacking MutH. In these organisms, MutL contains a latent endonuclease activity that introduces incisions used as an entry site for excision factors (Kadyrova and Kadyrov, 2016). This nicking activity is stimulated by interaction with MutS and/or replication processivity factors, the prokaryotic β clamp and the archaeal*/*eukaryotic proliferating cell nuclear antigen (PCNA) (Pluciennik et al., 2010, Shimada et al., 2013, Smith et al., 2015). Association with asymmetrically-loaded processivity factors also plays a role in directing the MutL incision to the nicked (nascent) strand (Pluciennik et al., 2010). Moreover, transient nicks associated with DNA synthesis or introduced during removal of misincorporated ribonucleotides from DNA could be used as starting points of excision and strand discrimination signals (Ghodgaonkar et al., 2013, Lujan et al., 2013). In the final repair steps, a DNA polymerase resynthesizes the resulting excision gap and a DNA ligase seals any remaining strand nicks.

Essential features of MutS for effective MMR initiation are its large conformational changes upon binding to mismatches and nucleotides, and a robust allosteric signaling between ATPase and DNA binding sites (Groothuizen and Sixma, 2016, Hingorani, 2016). Crystal structures of the MutS dimer reveal a modular protein arranged in a disk-like shape with two adjacent large channels (Lamers et al., 2000, Obmolova et al., 2000, Warren et al., 2007). In the absence of DNA, ATP-bound MutS forms a closed clamp that renders MutS unable to load on DNA. Following ATP hydrolysis, clamp domains are in a dynamic open/close equilibrium that allows DNA to enter the upper channel of the MutS dimer (Bhairosing-Kok et al., 2019, Mendillo et al., 2010, Obmolova et al., 2000). Loading of ADP-bound MutS onto double-stranded DNA stabilizes a closed state of clamp domains while DNA binding domains adopt a relatively open conformation that allows searching for mismatches (Obmolova et al., 2000, Qiu et al., 2012). Upon mismatch recognition, DNA binding domains move toward each other allowing specific contacts of the highly conserved Phe and Glu residues with mismatched bases (Cho et al., 2012, Gradia et al., 1999, Lamers et al., 2000, Obmolova et al., 2000, Qiu et al., 2012, Sharma et al., 2013, Warren et al., 2007). These specific contacts induce rapid ADP dissociation, switching MutS to an ATP-bound state that adopts a sliding clamp conformation. In this state, clamp domains cross each other and mismatch binding and connector domains move away from the mismatch, pushing the DNA downward into a novel channel and exposing a binding site for MutL (Groothuizen et al., 2015). Notably, this ATP-bound MutS sliding clamp is critical for MMR as it communicates mismatch recognition to downstream MMR components involved in the following steps.

In this report, we demonstrate that MutS from *Pseudomonas aeruginosa* associated with primed DNA replication structures by interacting with a second DNA binding site. Recognition of mismatches within primed DNA induced a structural rearrangement of MutS, leading to a more compact conformation, and hampered the transition into a sliding clamp. These results reveal that MutS mismatch detection in DNA replication structures involves a different DNA binding mode, conformational change and intramolecular signaling from those critical for MMR initiation.

## RESULTS

### Replication DNA substrates are recognized by MutS

Biochemical studies have proved the extraordinary capacity of MutS proteins to recognize a broad range of DNA substrates. The repair factor can bind to mispairs and deletion/insertion loops as well as damaged bases within MMR double-stranded DNA substrates (dsDNA). MutS is also able to associate with other DNA structures like D-loops, four-stranded Holliday junctions and G quadruplexes (Ehrat et al., 2012, Honda et al., 2014, Snowden et al., 2004). However, and to our knowledge, no studies have performed a detailed evaluation of MutS interaction with primed replication DNA structures (pDNA). With this aim, MutS from *P. aeruginosa* was purified and its ability to bind to replication and MMR DNA substrates (Figure 1A) was tested by electrophoretic mobility-shift assays (EMSA) and surface plasmon resonance (SPR).

**Figure 1.**
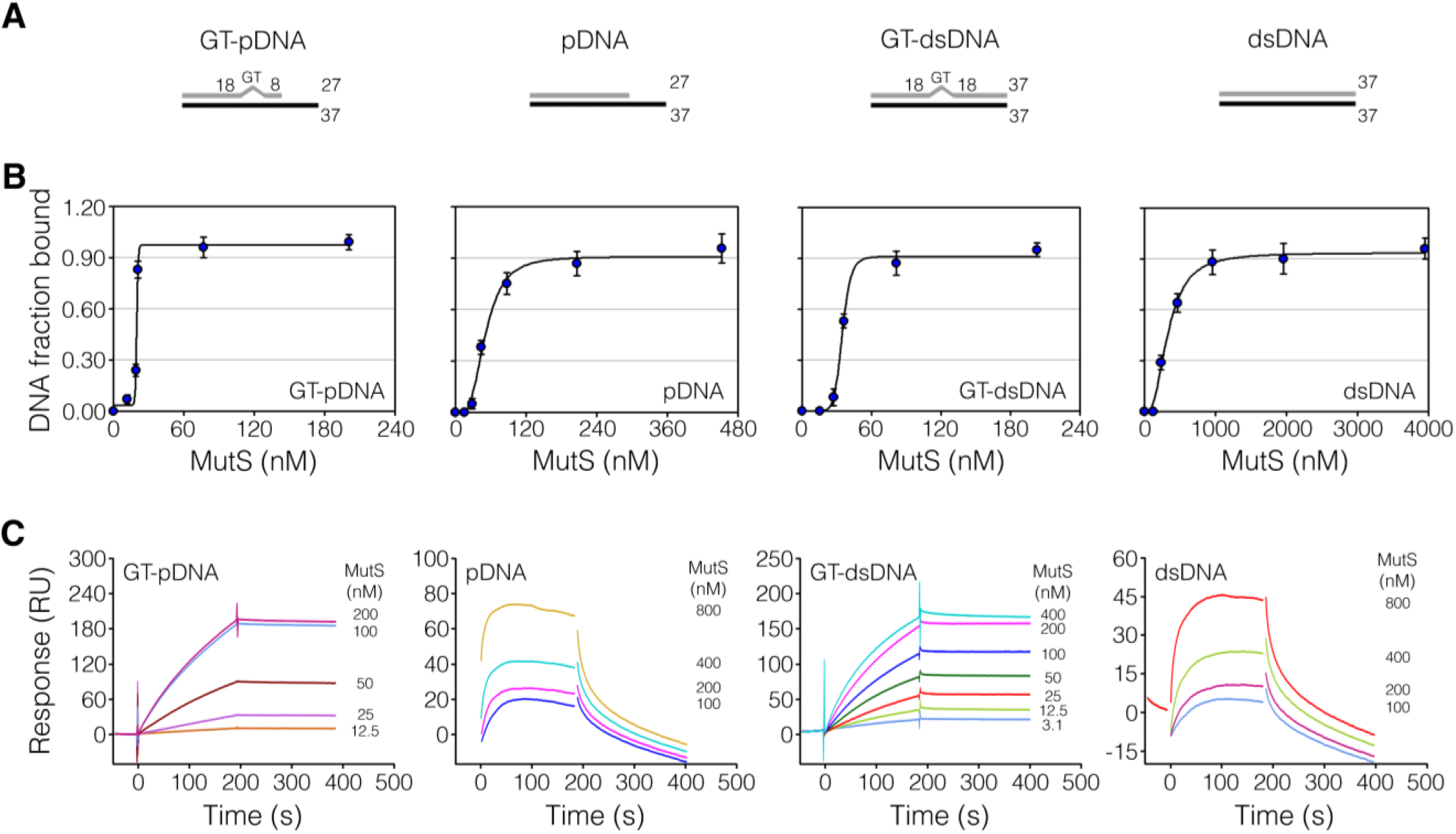
MutS association with replication DNA substrates. (A) Representation of the DNA substrates used in the present study. These substrates included a matched pDNA, conformed by a 27-nt primer strand and a 37-nt template strand, and a GT mismatched pDNA (GT-pDNA), which contained a GT mismatch 8 nt from the 3’-hydroxyl (OH) terminus. A dsDNA (37nt/37nt) and a GT-dsDNA, containing the mismatch centrally located, were used as controls. Numbers indicate the nucleotides at each strand and upstream and downstream of the mismatch. (B) Analysis of MutS interaction with DNA substrates by EMSA. MutS (15-4000 nM) was incubated with a fixed concentration of biotinylated DNA oligonucleotides (10 nM), and reaction products were analyzed on native polyacrylamide gels. Free and bound DNAs were detected by staining gels with infrared dye-labeled streptavidin. Binding curves were obtained by plotting the fraction of DNA bound versus free concentrations of MutS. Data represent mean ± standard deviation (n=3). (C) SPR measurements of MutS binding to biotinylated DNA substrates immobilized on a streptavidin-derivatized sensor chip (50-125 RU). Different protein concentrations (3-800 nM) are represented by different colors. Sensorgrams of MutS display solvent correction. Representative reference-subtracted curves are shown (n=3).

MutS was able to bind to all DNA substrates in EMSA assays (Figure 1B, and Supplementary Figure S1A). The plot of the DNA bound fraction as a function of free MutS concentrations fitted to a sigmoidal curve for all DNAs, indicating a cooperative interaction independently of the type of DNA structure. MutS displayed a strong interaction with GT-pDNA (*K*_D_= 20.0±0.1 nM) while its binding affinity was slightly weaker for pDNA (*K*_D_= 50.7±2.3 nM). As expected from previous studies, MutS exhibited a high affinity for GT-dsDNA (*K*_D_= 35.0±0.9 nM) whereas it bound to dsDNA with low efficiency (*K*_D_= 330.8±15.9 nM).

The SPR measurements showed that MutS specifically bound to GT-pDNA, exhibiting a K_D_ of 53.3±3.7 nM (Figure 1C and Supplementary Figure S1B). A cooperative interaction was observed as the plot of the maximum response units against MutS concentrations displayed a sigmoidal curve. MutS bound to GT-dsDNA with a K_D_ of 118.2±8.9 nM (Figure 1C and Supplementary Figure S1B). Binding data revealed a hyperbolic binding curve, which suggests that MutS exhibits a unique site of interaction for GT-dsDNA in our SPR conditions. It also indicates that MutS did not interact with two immobilized DNA strands within the range of DNA immobilization levels used in these experiments. Taking this into account, it is possible to hypothesize that MutS interaction with GT-pDNA involves two binding sites on a single immobilized DNA strand. For both mismatched DNA substrates, fitting of the kinetic data to a model, like a global 1:1 Langmuir model, was poor (Supplementary Figure S1C). Kinetics rates were outside the limits of the instrument and thus *K*_D_ values could not be accurately calculated. MutS associated with pDNA and dsDNA, albeit with a lower affinity (Figure 1C and Supplementary Figure S1C). The kinetic data for these DNA substrates fitted to a global 1:1 Langmuir model, which allowed the calculation of the kinetic parameters and the K_D_. MutS showed a K_D_ of 367 nM for pDNA (*k*on=1.2±0.1 10^5^ M^-1^ s^-1^, *k*off=4.4±0.2 10^-2^ s^-1^) and 1957 nM for dsDNA (*k*on=2.3±0.6 10^4^ M^-1^ s^-1^, *k*off=4.5±0.3 10^-2^ s^-1^). Taken together, our findings indicate that MutS specifically binds to replication DNA structures, containing or not a mismatch.

### Replication DNA substrates containing a mismatch alters MutS electrophoretic migration under native conditions

We next analyzed the mobility of the MutS-DNA complexes on native gels, which could provide information on their size or conformation. MutS was incubated in the absence or presence of DNA substrates and then, subjected to electrophoretic analysis. Gels were stained with ethidium bromide and subsequently with coomassie blue to detect DNA and protein, respectively, in all assays. We identified MutS-DNA complexes as bands that showed both ethidium bromide and coomassie blue staining (Figure 2A and Supplementary Figure S2A). Coomassie-stained gels are shown from now on for simplicity. In the absence of DNA, MutS migrated as a single band corresponding to a molecular weight of ∼448 kDa (band b), indicating that the protein is mainly a tetramer since the estimated molecular mass of the MutS monomer is 95 kDa. MutS bound to GT-pDNA exhibited a significant shift in mobility. ∼90% of the protein migrated as a band near ∼331 kDa (band a), which could be due to a change in the oligomeric state from tetramer to dimer or a conformational rearrangement to a more compact protein structure. This MutS mobility shift was only observed in the presence of at least a 2-fold excess of GT-pDNA (Supplementary Figure S2B), suggesting that a single MutS tetramer must bind to two GT-pDNA molecules in order to increase its mobility. Conversely, MutS complexed with GT-dsDNA showed a decreased mobility as ∼92% of the protein-DNA complex migrated at ∼694 kDa (band c). dsDNA (∼631 kDa) and pDNA (∼685 kDa) induced a similar effect on MutS migration to that observed for GT-dsDNA. This decreased migration could be due to a more extended conformation adopted by the MutS tetramer upon binding to these DNA substrates or binding of multiple tetramers on a single DNA molecule.

**Figure 2.**
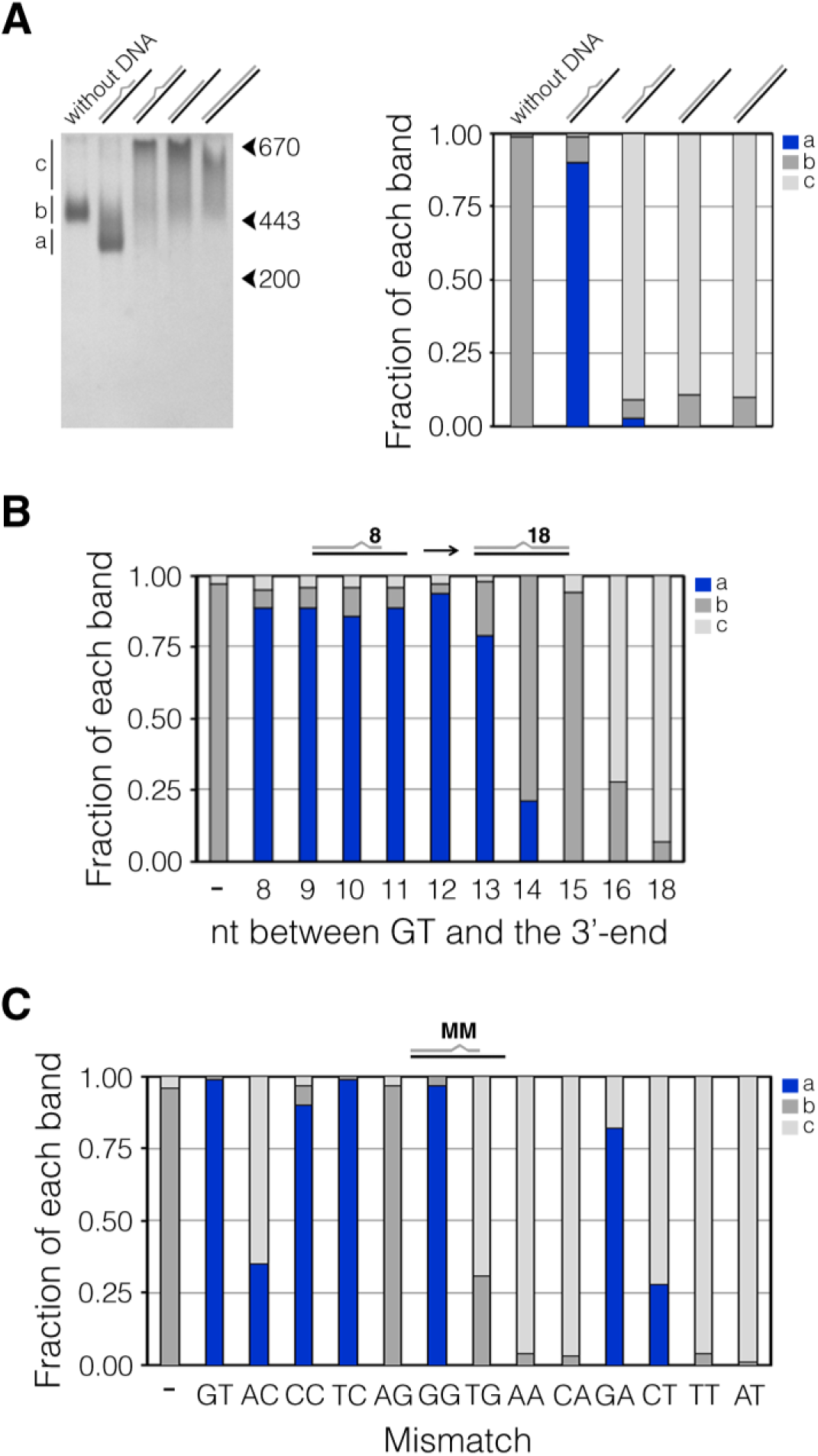
Electrophoretic migration of DNA-bound MutS under native conditions. MutS (2 μM) was individually incubated alone or with DNA substrates (4 μM). Reaction mixtures were analyzed on native polyacrylamide gels. (A, left panel) Representative image of a coomassie blue-stained gel showing the MutS-DNA mobility shift relative to free MutS. Migration of molecular weight markers, thyroglobulin (670 kDa), apoferritin (443 kDa) and β-amylase (200 kDa), is shown with arrows. a, b and c corresponds to the migration of MutS-GT-pDNA, free MutS, and MutS complexed with pDNA, GT-dsDNA or dsDNA bands, respectively. (A, right panel) Fraction of a, b and c bands for MutS bound to the different DNA structures. The intensity of a, b and c bands was quantified in each line to calculate the relative fraction of each band. (B) Fraction of a, b and c bands for MutS bound to GT-pDNA structures containing 8 nt to 16 nt between GT and the 3’-OH terminus of the primer strand. Control reactions without DNA (-) and GT-dsDNA (18 nt from the GT) were included. (C) Fraction of a, b and c bands for MutS bound to pDNA structures containing the twelve different mismatches (MM) located 8 nt from the 3’-OH terminus of the primer strand. Control reactions without DNA (-) and pDNA (AT) were included. Data represented the means of at least triplicates. Standard deviations were <5% of the mean.

In order to gain insight into the structural DNA features responsible for inducing a faster mobility of MutS by GT-pDNA, we determined the effect of extending the 3’-OH terminus of the primer strand in GT-pDNA until obtain the GT-dsDNA structure (Figure 2B and Supplementary Figure S2C). For this, different DNA substrates containing the GT mismatch located 8 nt (as in GT-pDNA) to 18 nt (as in GT-dsDNA) from the 3’-end were generated by adding 1 nt at a time at the 3’-OH terminus of the primer strand. Only GT-pDNAs with the GT mismatch located up to 13 nt far from the 3’-end induced a faster mobility of MutS (79-94% of protein in band a). MutS bound to the other GT-pDNAs exhibited a mobility similar to the DNA-unbound MutS or the MutS-GT-dsDNA complex. We also examined MutS migration on native gels in the presence of pDNAs containing all possible 12 mismatches (Figure 2C and Supplementary Figure S2D). Binding to pDNAs with CC, TC, GG and GA mismatches induced a faster mobility of MutS (82-99% of protein in band a). AC and CT mismatches produced a similar effect on MutS mobility, but to a lesser extent (35% and 28% of protein in band a, respectively). MutS bound to the other mismatches exhibited a migration similar to that observed for the matched pDNA (TG, AA, CA and TT) or free protein (AG).

To sum up, these data demonstrate that MutS binding to GT-pDNA provokes a faster migration of the protein relative to the DNA-unbound MutS on native gels. To induce this effect, the mismatched pDNA have to fulfill some important structural characteristics: a maximum separation of 13 nt between the mismatch and the 3’-OH terminus in the primer strand and the presence of a specific type of mismatch. On the other hand, GT-dsDNA, dsDNA and pDNA induce a retarded migration of the repair protein.

### MutS binding to mismatched replication DNA substrates induces a rearrangement of the protein structure

Native electrophoretic mobility assays suggested that MutS undergoes a conformational change upon binding to mismatched pDNA structures, which could result in a tetramer to dimer transition or in a more compact protein structure. To determine if the interaction with GT-pDNA triggers a change in the oligomeric state of MutS, two dimeric MutS mutant versions, MutSR842E and MutSΔ798, were analyzed (Miguel et al., 2008). Native gel analysis revealed that the MutS-GT-pDNA complex (∼331 kDa) migrated slower than DNA-unbound full-length MutSR842E (∼193 kDa) (Supplementary Figure S3A). Moreover, MutSR842E bound to GT-pDNA showed an increased mobility (∼136 kDa, band a) relative to the uncomplexed protein (Figure 3A). Association with GT-dsDNA, dsDNA or pDNA reduced the migration of MutSR842E (Figure 3A). Most of the DNA-complexed MutSR842E migrated at ∼316, 425 and 388-kDa, respectively (band b). As observed for the wild type protein, GT-pDNA induced a fast mobility of MutSR842E when the GT mismatch was located up to 13 nt from the 3’-OH terminus (Figure 3B and Supplementary Figure S3B). Furthermore, pDNA containing AC, CC, TC, GG, TG, CT, TT and GA triggered a faster mobility of MutSR842E, albeit to a lesser extent than GT (Figure 3C and Supplementary Figure S3C). The truncated MutSΔ798 mutant bound to GT-pDNA also exhibited an increased mobility relative to the DNA-unbound protein whereas interaction with GT-dsDNA, dsDNA and pDNA slightly retarded its migration (Supplementary Figure S3A). These results clearly indicate that GT-pDNA has no effect on the oligomeric state of MutS and both the MutS tetramer and dimer display a fast mobility upon binding to GT-pDNA.

**Figure 3.**
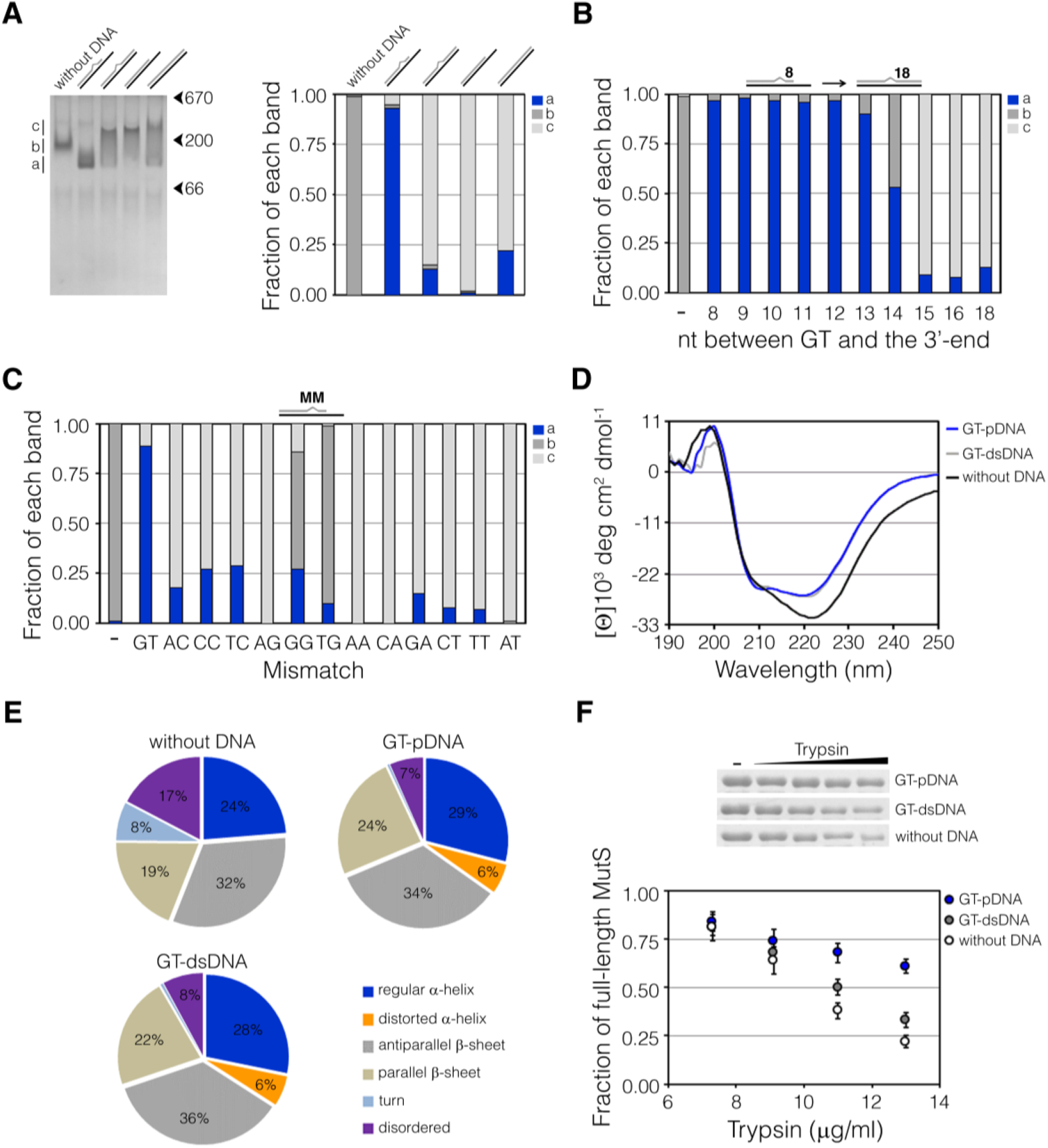
Rearrangement of the MutS structure upon binding to mismatched replication DNA. (A-C) Electrophoretic migration of MutSR842E bound to DNA under native conditions. MutSR842E (4 μM) was individually mixed with DNA substrates (4 μM), and reaction products were analyzed on native polyacrylamide gels. (A, left panel) Representative image of a coomassie blue-stained gel showing the MutSR842E-DNA mobility shift relative to the apo MutSR842E form. Thyroglobulin (670 kDa), β-amylase (200 kDa) and bovine albumin (66 kDa) band migration is shown with arrows. a, b and c corresponds to the migration of MutSR842E-GT-pDNA, free MutSR842E, and MutSR842E complexed with pDNA, GT-dsDNA or dsDNA bands, respectively. (A, right panel) Fraction of a, b and c bands for MutSR842E bound to DNA structures. The intensity of a, b and c bands was quantified in each line to calculate the relative fraction of each band. (B) Fraction of a, b and c bands for MutSR842E bound to GT-pDNA structures containing 8 nt to 16 nt from the GT to the 3’-OH terminus of the primer strand. Control reactions without DNA (-) and GT-dsDNA (18 nt from the GT) were included. (C) Fraction of a, b and c bands for MutSR842E bound to pDNA structures containing the twelve different mismatches (MM) located 8 nt from the 3’-OH terminus of the primer strand. Control reactions without DNA (-) and pDNA (AT) were included. Data represented the means of at least triplicates. Standard deviations were <5% of the mean. (D) Far-UV CD spectra of 2 μM MutS in the absence (black line) or presence of 4 μM GT-pDNA (blue line) and GT-dsDNA (grey line). The plotted spectra are the average of three independent scans. Contributions of buffer and DNA to ellipticity were subtracted. (E) Fraction of secondary structures estimated from CD spectra using the BeStSel web tool. (F) Limited proteolysis. 2 μM MutS was preincubated alone or with DNA substrates (4 µM), and then subjected to partial tryptic digestion. Reactions were analyzed by SDS-PAGE, and the intensity of full-length MutS bands was quantified and used to calculate the fraction of MutS remaining. The intensity of the MutS band in the control condition without trypsin was taken as 1. Data represent mean ± standard deviation (n=3).

We next examined if the binding to GT-pDNA changed the secondary and/or tertiary structures of MutS using far-UV circular dichroism (CD) and limited trypsin proteolysis assays, respectively. The CD spectra of MutS in the absence or presence of DNA were recorded (Figure 3D and Supplementary Figure S3D), and used to estimate the fraction of secondary structures by the BeStSel web tool (Micsonai et al., 2018) (Figure 3E and Supplementary Figure S3E). MutS association with GT-pDNA reduced the disordered structure fraction and increased the α-helix content when compared with free MutS. The other DNA substrates produced a similar effect on the secondary structure of MutS to that observed for GT-pDNA. Hence, MutS binding to DNA substrates induces further ordering of the secondary protein structure in a similar manner. In limited proteolysis experiments (Figure 3F), free MutS was readily cleaved by trypsin as ∼22% of the full length protein was detected at the highest trypsin concentration used. This correlated with the high number (92) of predicted cleavage sites for trypsin displayed by MutS. DNA binding protected MutS from proteolysis. However, the MutS-GT-pDNA complex exhibited a higher resistance to proteolysis relative to MutS-GT-dsDNA as ∼61% and ∼33% of full length MutS was detected at the highest trypsin concentration, respectively. This result suggests that the mismatched replication substrate induces a rearrangement of the tertiary structure of MutS to a more compact conformation, which is less susceptible to tryptic digestion.

### MutS bound to replication DNA structures containing a mismatch is more resistant to ATP-dependent release

The sliding clamp state has been detected in biochemical assays. MutS slides off short mismatched double-stranded oligonucleotides while it remains bound to oligonucleotides with both blocked ends in the presence of ATP (Acharya et al., 2003, Blackwell et al., 2001, Gradia et al., 1999). To test whether ATP induces MutS dissociation from replication DNA substrates, MutS was bound to DNA oligonucleotides with free ends and then, complexes were challenged with different ATP concentrations (Figure 4A and Supplementary Figure S4A). As previously described, the repair protein was released from GT-dsDNA upon the addition of ATP. The effective concentration of ATP for 50% MutS dissociation (EC_50_) was calculated to be 618±49 µM. ATP was less effective at inducing the dissociation of MutS from GT-pDNA, as the EC_50_ was 1182±68 µM. The addition of ATP promoted the release of MutS from pDNA (EC_50_=344±31 µM) and dsDNA (EC_50_=496±43 µM) more efficiently than that observed for the mismatched DNA substrates. These results show that MutS bound to a mismatch in replication DNA structures exhibits an enhanced stability upon ATP challenge.

**Figure 4.**
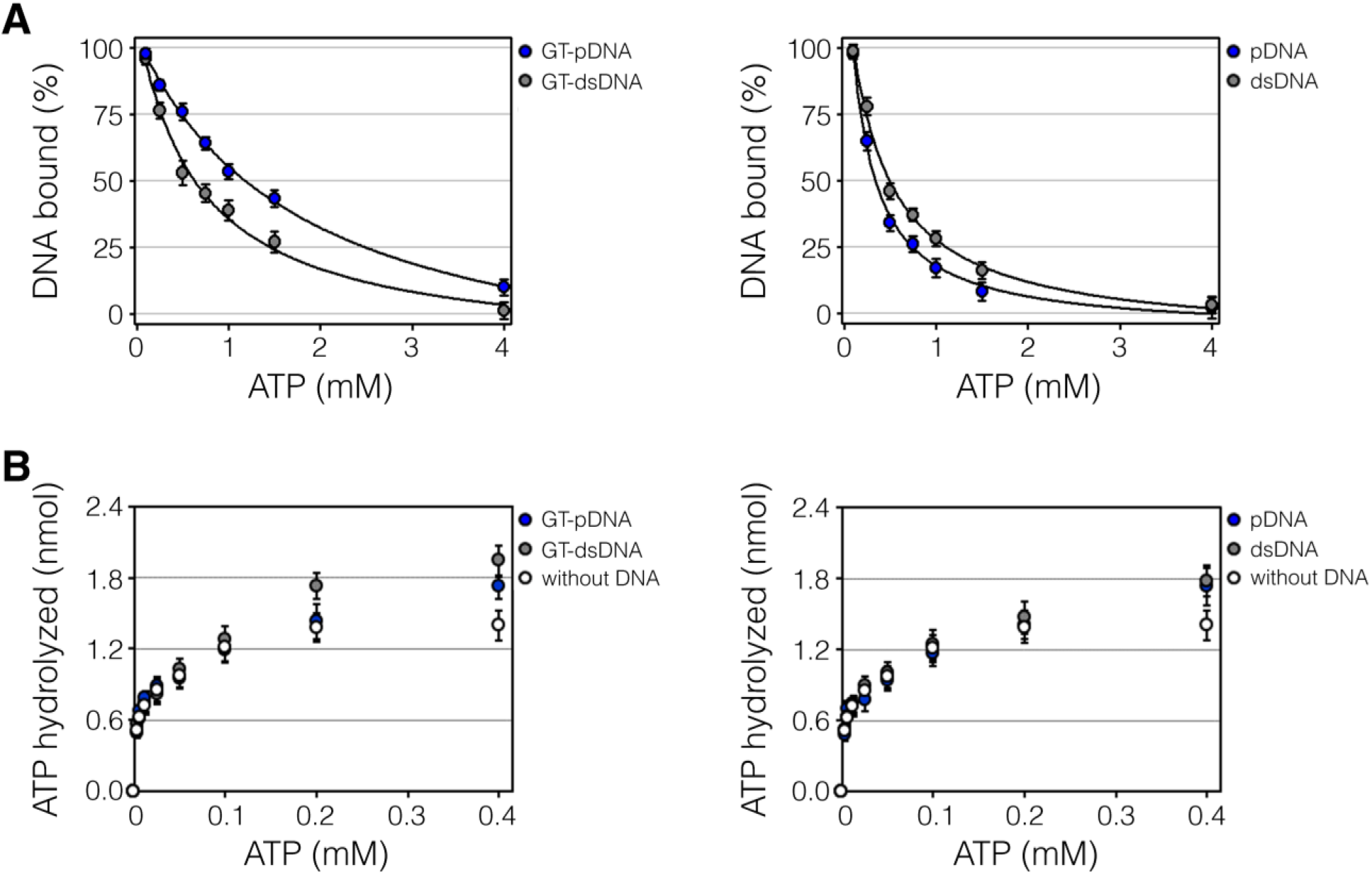
MutS association with replication DNA structures upon ATP challenge. (A) ATP-dependent release of MutS prebound to DNA substrates. MutS (250 nM) bound to DNA oligonucleotides (50 nM) was incubated with increasing concentrations of ATP (0.1-4.0 mM). Reaction products were analyzed on native polyacrylamide gels. Band intensities of free DNA were quantified and used to calculate the percentage of DNA bound. The EC50 was obtained by fitting curves as described in Material and Methods. (B) ATPase activity of MutS in the presence of DNA. ATP hydrolysis by MutS (40 nM) was measured in the absence or presence of DNA (1.6 µM) as a function of ATP concentration (3-400 µM). Data represent mean ± standard deviation (n=3).

It has been well documented that the ATPase activity of MutS is stimulated by a mismatch in dsDNAs with free ends as mismatch recognition stimulates ADP-ATP exchange and subsequent dissociation of ATP-bound MutS from DNA increases ATP hydrolysis (Blackwell et al., 2001, Gradia et al., 1999, Mendillo et al., 2005). According to our previous results, it can be predicted that GT-pDNA should be less efficient to promote the ATPase activity of MutS compared to GT-dsDNA. We investigated the effect of DNA substrates on the ATPase activity of MutS by measuring ATP hydrolysis in the absence or presence of a fixed concentration of DNA oligonucleotides with free ends and increasing ATP concentrations (Figure 4B). Hydrolysis of ATP by MutS was increased as ATP was titrated into the reaction. GT-dsDNA stimulated the ATPase activity between 200 and 400 µM ATP (∼1.4-fold). Conversely, GT-pDNA only increased this activity at 400 µM ATP (∼1.2-fold), exhibiting a similar behavior as matched DNA substrates. Both pDNA and dsDNA increased the ATP hydrolysis at the highest ATP concentration (∼1.2 and 1.3-fold). The experiments described above demonstrate that a mismatch within pDNA is less effective at stimulating the ATPase activity of MutS.

### MutS contains a novel putative DNA interaction site for DNA replication structures

We hypothesized that association of the repair factor with replication DNA substrates involves novel protein residues in the disk-like MutS protein (Figure 5A). The presence of a novel putative DNA binding site in MutS was first evaluated using the web server BindUP (Paz et al., 2016), which predicts nucleic acid binding residues based on the electrostatic features of the three-dimensional (3D) protein surface. There is no 3D-structure of *P. aeruginosa* MutS, therefore, we used crystal structures from *E. coli* MutS. Because of the high sequence identity (59%) between both proteins, it is expected that they share at least 80% of the structure (Miguel et al., 2008). Moreover, we determined that GT-pDNA also induced a higher mobility of *E. coli* MutS on native gels (Supplementary Figure S4B), which suggests that the conformational change induced by interaction with GT-pDNA is conserved between both bacterial MutS proteins. Using the high-resolution 3D-structure of the full-length dimeric MutSD835R mutant from *E. coli* solved in complex with a GT-dsDNA (Groothuizen et al., 2013), three positive electrostatic surface patches were predicted by BindUP (Figure 5B and Supplementary Data). One of the predicted positive patches included residues from the clamp and mismatch-binding domains, highly overlapping with the identified mismatch binding interface (referred here as DNA binding site I). The other predicted DNA binding surface consisted of amino acids from the ATPase, helix-turn-helix and C-terminal domains. Residues present in the ATPase domain included His 760, which stacks the adenine nucleotide, and the catalytic amino acids Asn 616 and His 728 (Lamers et al., 2000). It is possible that residues present in the ATPase domain were predicted to bind DNA because they are involved in the interaction with nucleotides. It is worthy to mention that no interaction was observed between the helix-turn-helix and C-terminal domains and DNA substrates in EMSA assays (Supplementary Figure S4C). Finally, a third positive surface including residues from the connector and core domains, which localizes in the second channel, was identified (referred here as DNA binding site II). This positive surface was also predicted to be a potential DNA binding site for crystal structures of all *E. coli* MutS, *Thermus aquaticus* MutS and human MutSα bound to a heteroduplex DNA (Supplementary Data).

**Figure 5.**
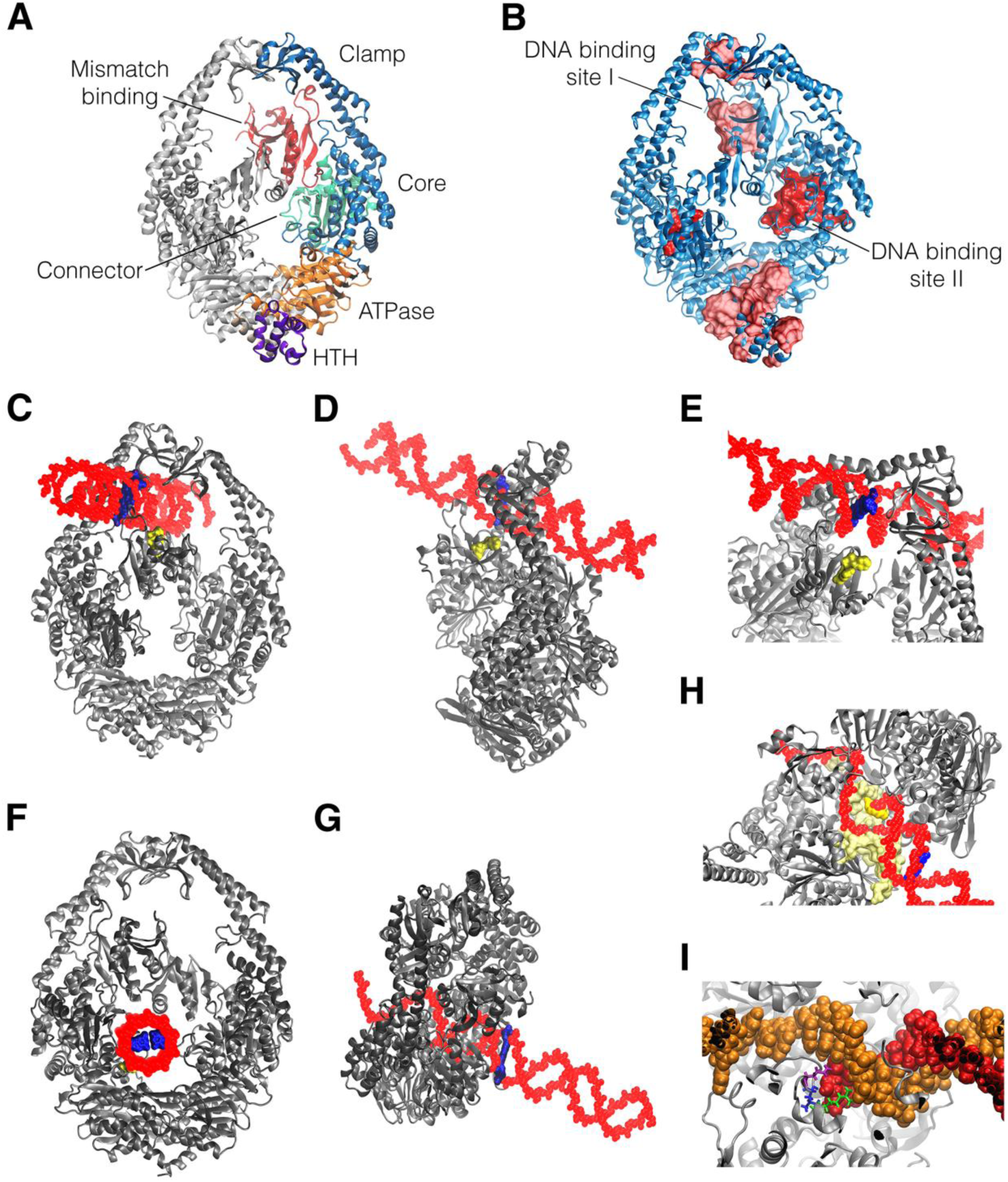
MutS contains a novel interaction site for replication DNA structures. (A) Ribbon representation of the *E. coli* MutSD835R crystal structure (PDB entry: 3ZLJ). One monomer is shown in gray and the other is coloured by domain. Mismatch binding domain (red), connector domain (green), core domain (light-blue), clamp domain (dark-blue), ATPase domain (orange) and helix-turn-helix domain (purple). The C-terminal domain is not shown for simplicity. (B) Putative DNA binding sites according to BindUp. Ribbon diagram of *E. coli* MutSD835R (PDB entry: 3ZLJ, blue) showing predicted DNA binding residues (surface representation, red). One of the sites was in the upper channel of MutS (light-red, referred here as DNA binding site I). Other of the putative binding sites was located in the ATPase and helix-turn-helix domains (light-red). The lower channel shows the putative binding site referred here on as DNA binding site II (dark-red). (C-I) Docking simulations of the *E. coli* MutSΔ800 (PDB entry: 1E3M) interaction with mismatched dsDNA and pDNA. MutSΔ800 is represented by ribbons diagrams (grey) and DNAs are showed in a space-filling model. Front (C) and lateral (D) view of the 3D structure of MutSΔ800 with docked GT-dsDNA. (E) Close view of DNA binding site I in MutSΔ800 bound to GT-dsDNA, showing the GT mismatch (blue) and the Phe 36 and Glu 38 residues (surface representation, yellow). Front (F) and side (G) view of the 3D docked structure of GT-pDNA to MutSΔ800. (H) Residues (surface representation, yellow) from the DNA binding site II contacted the double-stranded and single-stranded regions of the GT-pDNA. (I) Close view of the interaction between the 3’-OH terminus of GT-pDNA and Arg 275, Gln 281 and Asn 282 residues (stick representation; green, blue and magenta, respectively) in the DNA binding site II. The primer and template strands are shown in red and orange, respectively.

We next performed molecular docking of the interaction between the *E. coli* MutSΔ800 crystal structure (Lamers et al., 2000) and the modelled 3D-structure of DNA substrates. Docking of GT-dsDNA to the *E. coli* MutSΔ800 structure showed that the most frequent DNA-MutS complex consisted in the GT-dsDNA bound to the DNA binding site I (Figure 5C to 5E, and Supplementary Table S1). On the other hand, most of the docked *E. coli* MutSΔ800-GT-pDNA structures showed that GT-pDNA interacted with the DNA binding site II, contacting residues present in the connector and core domains (Figure 5F to 5H, and Supplementary Table S1). All these residues always belonged to one of the two MutS monomers, similarly to the asymmetry in mismatch binding at the DNA binding site I. The pDNA structure exclusively docked in the DNA binding site II while dsDNA was predicted to bind to both DNA binding sites (Supplementary Figures S5A and S5B, and Table S1). Finally, a single-stranded DNA, corresponding to the template strand, was not capable of binding to neither DNA binding site I nor site II (Supplementary Figure S5C and Table S1).

To confirm the interaction of GT-pDNA to the DNA binding site II, we performed molecular dynamics (MD) simulations using the docked structure of MutS-GT-pDNA as starting structure. Root mean square deviations (RMSD) of MutS and DNA backbone chains were analyzed to determine the evolution and stability of the system (Supplementary Figure S6A). RMSD values were stable for MutS (3-4 Å) and DNA (4-6 Å for the primer strand and 6-9 Å for the template strand) after 20 ns and 10 ns, respectively. GT-pDNA remained bound to MutS along the MD simulation, indicating that this is a stable interaction. We next calculated averages minimum distances between both MutS monomers (named chain A and chain B) and the primer and template strands from MD simulations (Supplementary Figure S6B). This analysis, along with visual inspection of the simulated end-structure, allowed us to determine which MutS residues are in close contact with GT-pDNA. The single-stranded DNA region of GT-pDNA interacted with the A chain of MutS, specifically with two loops encompassing residues Glu 278-Thr 290 and Gly 583-Ala 598, and the alpha helix formed by residues Thr 335-Val 345. Also, a second helix in chain B (residues Tyr 41 to Asp 52) was involved in this interaction. The double-stranded region of GT-pDNA interacted with an alpha helix (Ala 181-Arg 198) in chain A and loops Thr 280-Thr 290 and Val 591-Phe 596 from chain B. Notable, the connector Arg 275, Gln 281 and Asn 282 residues were in close proximity to the 3’-OH terminus (Figure 5I).

Finally, hydrogen bonds formed between MutS and GT-pDNA during the course of simulation were analyzed. This type of bond increased during MD, when compared to the initial docked structure, reaching an average value of 26 bonds formed (Supplementary Figure S6C). Residues involved in hydrogen bonds are depicted in Supplementary Table S2. Confirming the key role of Arg 275, Gln 281 and Asn 282 in the interaction of MutS with primed DNA, these protein residues formed stable hydrogen bonds (Supplementary Figure S6D).

To sum up, our bioinformatics data provide support for a novel DNA binding surface located in the second channel of MutS, which may be involved in the specific association with replication DNA structures. Within this new binding site, Arg 275, Gln 281 and Asn 282 residues are predicted to be implicated in the direct interaction with the 3’-OH terminus. Arg 275 and Asn 282, but not Gln 281, are conserved in *P. aeruginosa* MutS, corresponding to Arg 272 and Asn 279.

### Mutation of Asn 279 and Arg 272 in MutS alters binding to replication DNA structures

Based on our data, Asn 279 and Arg 272 in *P. aeruginosa* MutS could be implicated in the interaction with replication DNA structures. To address the role of these residues in the function of MutS, we carried out the biochemical analysis of the MutSN279A, MutSR272A and MutSR272E mutants. We also characterized MutSF33A, a mutant version of the mismatch binding Phe located at the DNA binding site I. It is important to note that MutSN279A, MutSR272A and MutSF33A exhibited almost the same ATPase activity as MutS, however, MutSR272E showed a significant decrease in the ATPase activity (Supplementary Figure S7A). All mutant versions displayed a similar affinity for the MutS-interacting partners, MutL and β clamp, compared to the wild-type protein (Supplementary Figure S7B and S7C). In addition, N279A, R272A and R272E and F33A mutations did not alter the quaternary structure of MutS as the mutant versions formed mainly tetramers as the wild type protein (Supplementary Figure S7D). These data indicate that N279A, R272A and F33A mutations have no effect on the ATPase activity, interaction with MutL and β clamp, and the oligomeric state of MutS. In contrast, the R272E change results in an important defect in the ATPase activity of MutS, and thus, MutSR272E was discarded from further analysis.

DNA binding activity of MutS mutants was assessed in EMSA assays (Figure 6A and 6B). MutSN279A exhibited a decreased interaction with GT-pDNA (*K*_D_= 63.2±2.4 nM) and pDNA (*K*_D_= 430.0±18.0 nM) compared to MutS, displaying 3- and 8-fold lower binding affinities, respectively. MutSR272A showed 2- and 3-fold reduced binding affinities for GT-pDNA (*K*_D_= 49.3±1.0 nM) and pDNA (*K*_D_= 143.2±8.6 nM), respectively. Binding affinities of both MutS mutants for double-stranded substrates were comparable to that observed for MutS. *K*_D_ values for binding to GT-dsDNA and dsDNA were estimated to be 41.3±5.2 and 358.3±36.9 nM for MutSN279A, and 38.2±0.7 and 308.7±20.9 nM for MutSR272A. A residual DNA binding was noticeable between the mismatch binding site I mutant, MutSF33A, and DNA substrates under the EMSA conditions (Supplementary Figure S7E). We also evaluated the mobility of MutSN279A and MutSR272A complexed with DNA on native gels (Figure S6C). Replication DNA substrates induced a different migration of both mutants when compared with MutS. MutSN279A and MutSR272A bound to pDNA (∼481 and 496 kDa, band b) migrated as the DNA-unbound proteins (∼452 and 467 kDa, band b), instead of showing a diminished migration like the MutS-pDNA complex (∼685 kDa). Interaction of MutS mutant versions with GT-pDNA increased the mobility, however, most of the protein migrated as a broad band between ∼355-420 kDa (band a). This is different from that observed for MutS, which ran as a sharp band (∼331 kDa). Conversely, MutSN279A and MutSR272A bound to GT-dsDNA (∼713 and 684 kDa, band c) and dsDNA (∼657 and 681 kDa, band c) showed a decreased mobility similar to MutS (∼694 and 631 kDa). These data clearly demonstrate that N279A and R272A substitutions result in a specific defect of MutS in binding to primed DNA structures.

**Figure 6.**
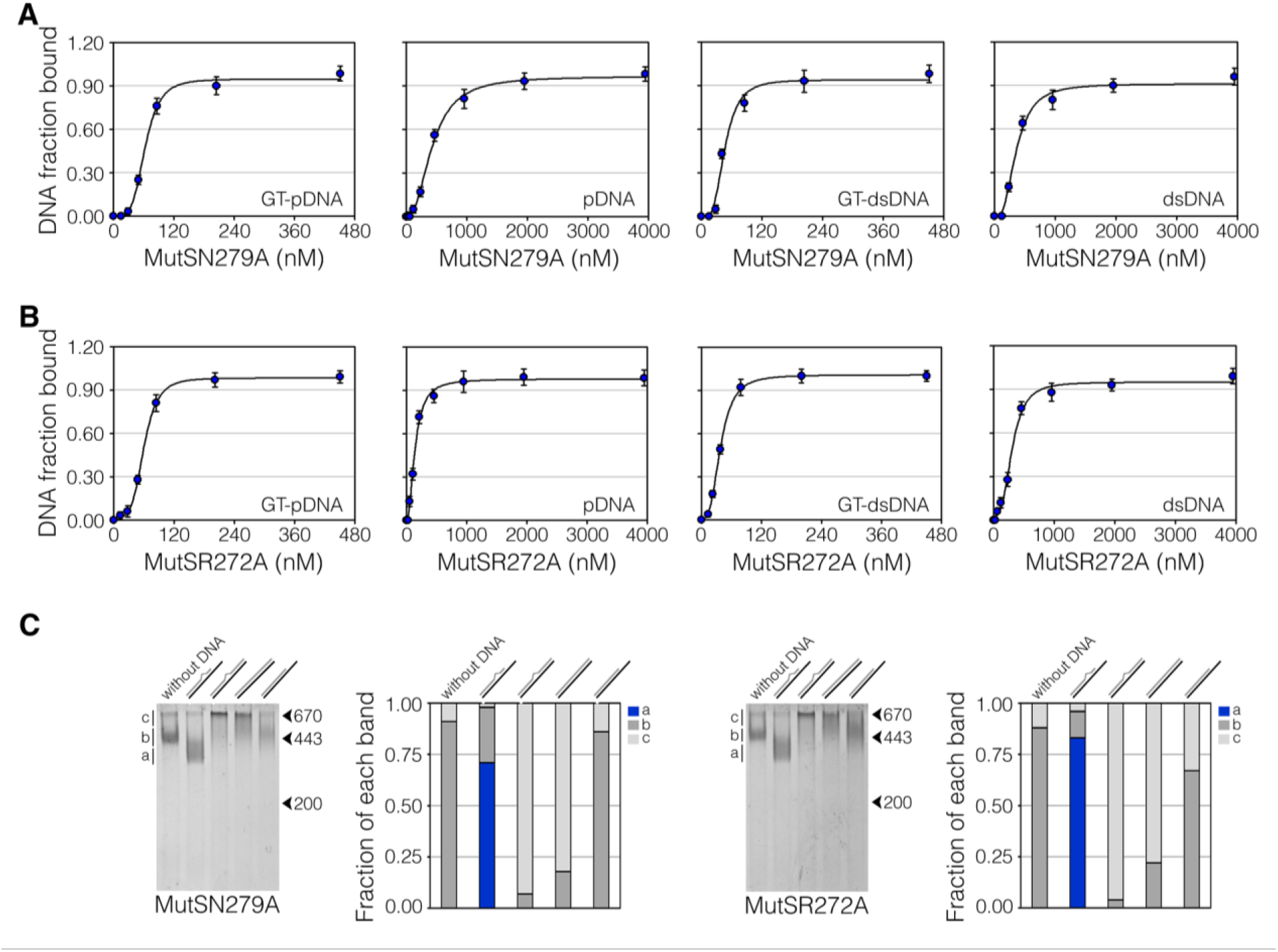
Effect of mutation of Asn 279 and Arg 272 on DNA binding of MutS. (A and B) Analysis of MutSN279A and MutSR272A interaction with DNA substrates by EMSA. MutS mutants (15-4000 nM) were incubated with a fixed concentration of biotinylated DNA oligonucleotides (10 nM). Reaction products were analyzed on native polyacrylamide gels and detected by staining with infrared dye-labeled streptavidin. Binding curves were obtained by plotting the fraction of DNA bound versus free concentrations of MutS mutants. Data represent mean ± standard deviation (n=3). (C) Electrophoretic migration of MutSN279A and MutSR272A bound to DNA under native conditions. MutS proteins (2 μM) were individually incubated alone or with DNA substrates (4 μM), and reaction mixtures were analyzed on native polyacrylamide gels. Representative images of coomassie blue-stained gels showing the mobility of MutS mutants bound to DNA relative to free proteins. Migration of molecular weight markers, thyroglobulin (670 kDa), apoferritin (443 kDa) and β-amylase (200 kDa), is shown with arrows. a, b and c corresponds to the migration of proteins bound to GT-pDNA, free proteins, and proteins complexed with pDNA, GT-dsDNA or dsDNA bands, respectively. The intensity of a, b and c bands was quantified in each line to calculate the relative fraction of each band. Data represented the means of at least triplicates. Standard deviations were <5% of the mean.

### MutSN279A and MutSR272A are proficient in mismatch repair *in vivo*

In order to analyze the *in vivo* antimutagenic activity of MutS mutants, we performed complementation assays by introducing pEx plasmids bearing the wild-type and mutant *mutS* alleles under the control of the IPTG inducible Ptac promoter into a *mutS*-deficient PAO1 strain. Expression of the ectopic *mutS* genes was detected by Western blot assays in cells harboring pEx plasmids after induction with IPTG (Supplementary Figure S8). Cellular levels of MutSN279A, MutSR272A and MutSF33A were similar to that of MutS, indicating that there are not significant differences in the expression of MutS proteins. MutS was not detected in IPTG-induced cells carrying the empty vector pEX.

Spontaneous mutations in different target genes on the *P. aeruginosa* chromosome were measured by estimating mutation rates to resistance to ciprofloxacin (Cip^r^), rifampicin (Rif^r^) and amikacin (Amk^r^) (Table 1). Cells expressing MutSN279A and MutSR272A showed mutation rates similar to that of transformants expressing MutS. In contrast, cells expressing MutSF33A displayed 35-, 54- and 19-fold higher mutation rates to Cip^r^, Rif^r^ and Amk^r^, respectively. Control cells harboring the empty plasmid exhibited 102-, 206 and 134-fold increased levels of mutations. These results show that MutSN279A and MutSR272A are capable of restoring mismatch repair at levels comparable to that of MutS. Conversely, F33A mutation results in a significant loss of mismatch repair capability *in vivo*.

**Table 1.**
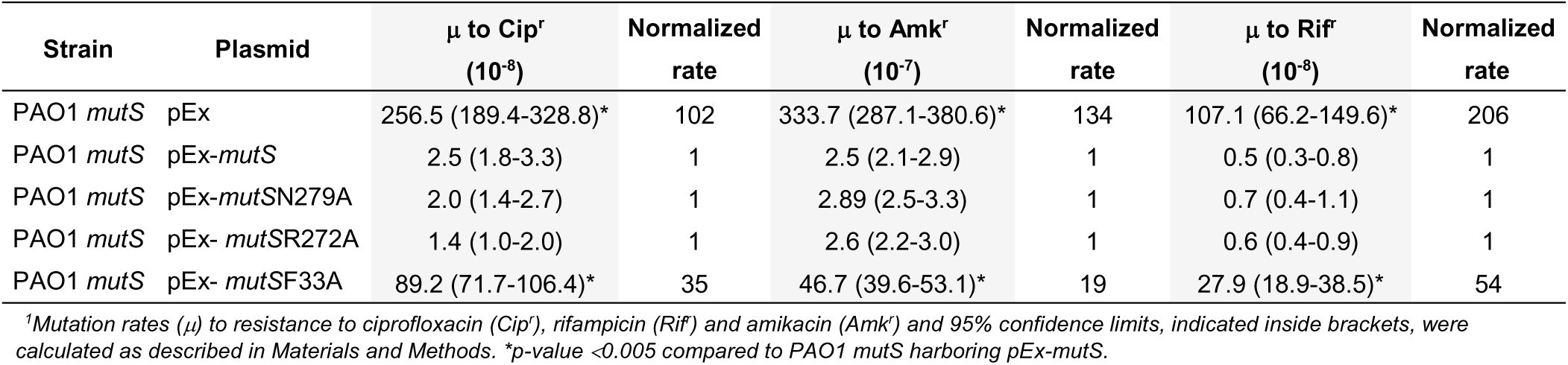
Complementation assays^1^

| Strain | Plasmid | $\mu$ to Cip<sup>r</sup><br>(10<sup>-8</sup>) | Normalized<br>rate | $\mu$ to Amk<sup>r</sup><br>(10<sup>-7</sup>) | Normalized<br>rate | $\mu$ to Rif<sup>r</sup><br>(10<sup>-8</sup>) | Normalized<br>rate |
| --- | --- | --- | --- | --- | --- | --- | --- |
| PAO1 *mutS* | pEx | 256.5 (189.4-328.8)* | 102 | 333.7 (287.1-380.6)* | 134 | 107.1 (66.2-149.6)* | 206 |
| PAO1 *mutS* | pEx-*mutS* | 2.5 (1.8-3.3) | 1 | 2.5 (2.1-2.9) | 1 | 0.5 (0.3-0.8) | 1 |
| PAO1 *mutS* | pEx-*mutSN279A* | 2.0 (1.4-2.7) | 1 | 2.89 (2.5-3.3) | 1 | 0.7 (0.4-1.1) | 1 |
| PAO1 *mutS* | pEx- *mutSR272A* | 1.4 (1.0-2.0) | 1 | 2.6 (2.2-3.0) | 1 | 0.6 (0.4-0.9) | 1 |
| PAO1 *mutS* | pEx- *mutSF33A* | 89.2 (71.7-106.4)* | 35 | 46.7 (39.6-53.1)* | 19 | 27.9 (18.9-38.5)* | 54 |

<sup>1</sup>Mutation rates ( $\mu$ ) to resistance to ciprofloxacin (Cip<sup>r</sup>), rifampicin (Rif<sup>r</sup>) and amikacin (Amk<sup>r</sup>) and 95% confidence limits, indicated inside brackets, were calculated as described in Materials and Methods. \*p-value <0.005 compared to PAO1 *mutS* harboring pEx-*mutS*.

## DISCUSSION

Most of the biochemical studies characterizing MutS activities have been carried out with double-stranded DNA (dsDNA), the substrate of the mismatch repair process. In the current work, we performed a detailed analysis of the interaction of MutS with primed DNA (pDNA) substrates and their effect on the different activities displayed by the repair factor. This kind of DNA structure is mainly present at and accumulates behind replication forks during lagging-strand synthesis (Moolman et al., 2014, Su’etsugu and Errington, 2011). Thus, we refer to it here as replication DNA substrate, albeit primed DNA can be an intermediate in repair and recombination processes.

Our data demonstrated that MutS from *P. aeruginosa* is able to interact with replication DNA structures, containing or not a mismatched base. Binding affinities of MutS to mismatched and matched pDNA were comparable to that observed for the mismatched dsDNA, following the order: GT-pDNA > GT-dsDNA > pDNA > dsDNA. Notably, MutS associated with pDNA with a similar affinity as replication factors involved in the recognition of primer-template junctions during DNA synthesis, the β clamp processivity factor and the clamp loader (Georgescu et al., 2008, Simonetta et al., 2009). In addition, our bioinformatics analysis predicted a novel DNA interaction surface (DNA binding site II) for replication DNA structures, which encompassed residues from the connector and core domains positioned in the lower channel of MutS. Among these residues, Asn 279 and Arg 272 in *P. aeruginosa* MutS appeared to directly interact with the 3’-OH terminus of pDNA by hydrogen bonding. Mutating these amino acids to Ala (MutSN279A and MutSR272A mutants) conferred a specific defect in the interaction of MutS with replication DNA substrates. Both mutant versions exhibited a significant decrease in binding affinity for pDNA, and to a lesser extent for GT-pDNA, compared to the wild-type protein. MutSN279A showed a major defect in binding to pDNA, which is in accordance with the fact that Asn 279 was predicted to form the most stable hydrogen bond. Conversely, N279A and R272A mutations did not impair mismatch recognition in dsDNA as association with GT-dsDNA was not affected. These findings clearly demonstrate that Asn 279 and Arg 272 correspond to novel DNA binding residues in *P. aeruginosa* MutS, which play an important role in the interaction with replication DNA structures.

It is important to note that the new DNA interaction site described in the present work is different from that observed in the sliding clamp conformation of *E. coli* MutS (Groothuizen et al., 2015). In this state, a large DNA binding channel is open up as *E. coli* MutS subunits tilt across each other and the connector and mismatch-binding domains are reoriented. Asn 282 and Arg 275 do not form part of this DNA interaction surface, as both residues do not line the large channel (Supplementary Figure S5D and E). These amino acids are located within a small channel at the bottom of *E. coli* MutS and are probably inaccessible for binding to the DNA helix in the sliding clamp conformation.

We also provide evidences of a different conformational change of MutS triggered by binding to mismatched pDNA. The repair protein appeared to adopt a more compact conformation by reorganizing its tertiary structure. This new state was revealed as a fast-migrating MutS-DNA complex on native gels, which exhibited a high resistance to proteolysis. We discarded that binding to GT-pDNA induces a specific rearrangement of the secondary structure or a change in the oligomeric state of MutS. The presence of GT-pDNA conferred MutS with a more ordered secondary structure, an effect produced by all evaluated DNA substrates. In addition, dimeric MutSR842E and MutSΔ798 mutants showed an increased mobility when bound to GT-pDNA, demonstrating that this DNA structure induces a protein compaction of the MutS dimer and tetramer. Understanding how MutS adopts this novel conformation remains to be elucidated. It is possible to postulate that the new MutS structure represents a state wherein both DNA binding sites are occupied in the dimer, the DNA binding site I by the mismatch and the DNA binding site II by the 3’-OH terminus. Both DNA interaction surfaces are distant as Arg 275 and Asn 282 are located ∼55 Å far from Phe 36 in the *E. coli* MutS structures (Supplementary Figure S5F-H). Thus, and taking into account that the distance between the mismatch and the 3’-OH terminus required to induce the MutS compaction was ∼27-44 Å (8-13 nt), DNA binding sites I and II should approach each other, resulting in a more compact MutS conformation. In this sense, our results suggest that mismatched pDNA interacts with both DNA binding sites in MutS. SPR data revealed a cooperative interaction of the repair factor with GT-pDNA, indicating two-binding sites for this DNA substrate. In contrast, a single binding site was shown for the other DNA substrates under SPR conditions. In addition, interaction with two DNA sites can explain the highly stable complex formed by MutS and mismatched pDNA. Dissociation of the repair factor from GT-pDNA was only accomplished by flowing a chaotropic agent over the SPR sensor surface. This harsh regeneration condition was not necessary to release MutS from immobilized GT-dsDNA, pDNA and dsDNA. Finally, interaction with the DNA binding site II is important for MutS binding to mismatched pDNA and the resulting conformational change. MutSN279A and MutSR272A displayed a decreased affinity by GT-pDNA and a different migration pattern to that observed for the wild-type protein in complex with this DNA.

A hallmark of the mismatch repair process is the ATP induced transition of MutS from a stationary clamp bound to a mismatch to a mobile sliding clamp (Acharya et al., 2003, Gradia et al., 1999, Groothuizen et al., 2015, Liu et al., 2016). Mismatch binding triggers ADP release and favors ATP binding. ATP-bound MutS changes into a conformation wherein mismatch binding domains move away from the mismatch, clamp domains cross each other and connector domains move outward. This transition allows MutS to adopt a mobile state, in which it remains because ATP hydrolysis is transiently suppressed. A similar molecular mechanism underlines MutS function in the inhibition of recombination between divergent sequences (Honda et al., 2014, Snowden et al., 2004). Our data indicated that MutS bound to a mismatch in replication DNA structures shows an enhanced stability upon challenge with ATP. This nucleotide was less effective in inducing the dissociation of MutS from GT-pDNA than GT-dsDNA. Accordingly, GT-pDNA was not as efficient as GT-dsDNA in stimulating the ATPase activity of MutS. In agreement with our findings, ATP is less efficient at triggering the release of human MutSα from mismatched pDNA than mismatched dsDNA structures contained in D-loops (D50+20^G/T^ vs D50^G/T^ DNA structures) (Honda et al., 2014). Interesting, ATP promoted the release of MutS from pDNA more efficiently than from mismatched pDNA. This result indicates that the unique DNA interaction with the DNA binding site II results in an efficient dissociation of MutS in the presence of ATP while the formation of a stable MutS-DNA complex requires DNA interaction with both DNA binding sites. One possibility to explain the enhanced stability of the MutS-GT-pDNA complex is that ADP-ATP exchange is less efficiently induced when the repair factor interacts with this DNA structure. MutS release from GT-pDNA required higher ATP concentrations, suggesting that ATP binding could be affected. Another possibility is that MutS can not effectively adopt the sliding clamp conformation upon mismatch recognition in pDNA. It is probable that, when both DNA binding sites are occupied in MutS, the ATP-induced reorientation of mismatch binding and connector domains is less likely to occur as both regions are simultaneously involved in DNA binding. In conclusion, MutS mismatch recognition in replication DNA structures induces a different signaling between DNA binding and ATPase sites from that triggered by binding to heteroduplex DNA.

Single amino acid substitution at Asn 279 and Arg 272 did not alter the repair function of MutS *in vivo*. MutSN279A and MutSR272A mutants were able to fully complement a *mutS* defect in *P. aeruginosa*. This correlates with the fact that N279A and R272A mutations had no effect on neither mismatch-binding nor other critical MutS activities for repair like ATP hydrolysis and MutL interaction. A similar result was shown for the Ala substitution of Arg 275 in *E. coli* MutS, as the resulting mutant version was as proficient as the wild-type protein in MMR (Junop et al., 2003). In contrast, and as it has been previously reported (Junop et al., 2003, Drotschmann et al., 2001), mutation of the mismatch-binding Phe residue (MutSF33A mutant) caused a noticeable defect in the *P. aeruginosa* MutS repair activity. Therefore, our findings establish that the novel DNA binding site II does no play a role in the correction of replication errors by MutS, which raises questions about the functional importance of MutS interaction with replication DNA structures.

Substantial evidence demonstrates a spatiotemporal coupling of MutS with DNA replication, which is thought to facilitate mismatch recognition during MMR (Haye and Gammie, 2015, Liao et al., 2015). MutS tracks with the replisome and persists transiently behind the advancing replication forks in living cells (Haye and Gammie, 2015, Liao et al., 2015). This localization of MutS at replication factories has been mainly ascribed to interactions with prokaryotic β clamp and eukaryotic PCNA (Haye and Gammie, 2015, Hombauer et al., 2011, Kleczkowska et al., 2001, Lenhart et al., 2013, Liao et al., 2015, Simmons et al., 2008). Our data revealed that MutS mismatch detection within pDNA involves a different DNA binding mode, conformational change and intramolecular signaling from that displayed during heteroduplex recognition (Figure 7). Accordingly, initiation of MMR could be impaired when MutS binds to mismatches within replication DNA structures as this factor can not elicit the canonical DNA repair pathway. This is consistent with the observation that active sites of repair do not colocalize with replisome components in cells (Schmidt and Hombauer, 2016). In addition, interruption of MutS association with processivity factors has a limited effect on MMR (Iyer et al., 2008, Pluciennik et al., 2009, Shell et al., 2007, Simmons et al., 2008). Thus, the different response triggered when MutS recognizes a mismatch within replication DNA structures could function as a mechanism to prevent MMR initiation at replication sites. This mechanism could be important for protecting cells from the detrimental effects of DNA replication disruption by MMR processing, as it has been described for the MMR processing of alkylation damage (Gupta et al., 2018). On the other hand, we reported that MutS contributes to control access of the low-fidelity DNA polymerase (Pol) IV to replication sites by regulating its association with β clamp (Margara et al., 2016). Our previous data showed that mismatched pDNA enhances MutS binding to β clamp and improves the ability of MutS to control Pol IV interaction with β clamp (Margara et al., 2016). Interestingly, the results presented here indicated that MutS compaction is strongly induced by GT, CC, TC, GG and GA in pDNA, which correspond to the preferred mismatches introduced by *P. aeruginosa* Pol IV (Margara et al., 2016). Therefore, it is tempting to speculate that detection of mismatches in replication DNA structures by MutS triggers control of Pol IV access to replication sites. Additional studies will be required to further establish these hypotheses.

**Figure 7.**
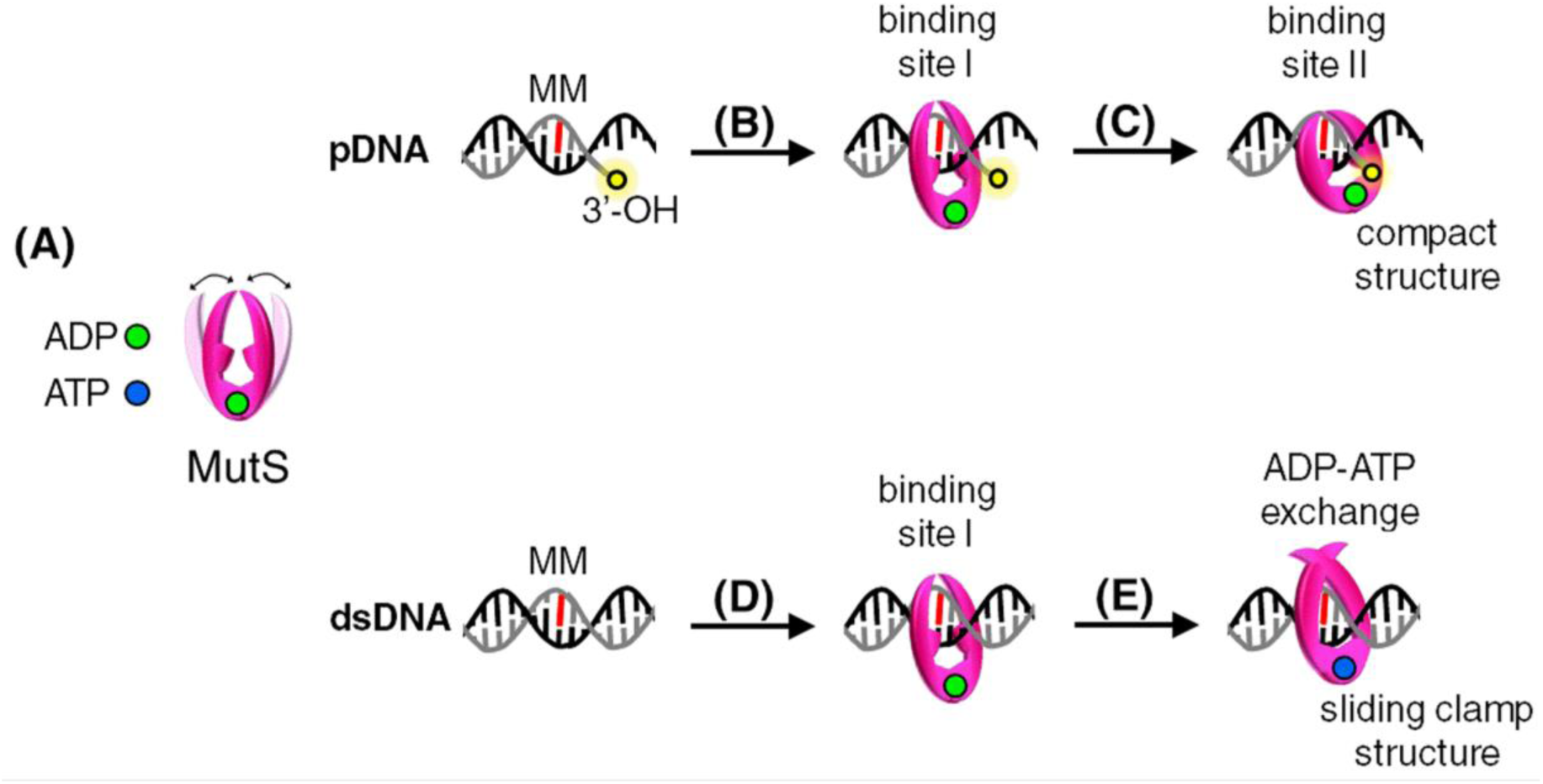
Proposed model for mismatch recognition within replication DNA structures by MutS. (A) Free-DNA MutS bound to ADP dynamically opens and closes its clamp domains, allowing MutS to load on DNA. (B) When ADP-bound MutS recognizes a mismatch (MM) located near to a primed site (pDNA), the repair factor closes its clamp domains and the Phe and Glu residues in the DNA binding site I contact the mismatch. (C) Then, MutS interacts with the 3’-OH terminus through the Asn and Arg residues in the DNA binding site II, leading to a conformational change to a more compact structure. Under this structural state, MutS is less prone to bind to ATP and can not effectively adopt a sliding clamp conformation. (D) When ADP-bound MutS detects a mismatch in double-stranded DNA (dsDNA), the DNA binding site I is occupied. (E) In the absence of a 3’-OH terminus, the DNA binding site II is free, which allows the exchange ADP for ATP and the conformational change into a sliding clamp.

## MATERIAL AND METHODS

### Bacterial strains, plasmids, oligonucleotides and chemicals

*P. aeruginosa* PAO1 *mutS* strain (Oliver et al., 2002) was provided by Dr. Oliver (Servicio de Microbiología, Hospital Son Dureta, Palma de Mallorca, Spain). *E. coli* Bl21 (λDE3) and DH5α strains were supplied by ThermoFisher Scientific. The expression plasmids pET15b and pMAL-c2X were obtained from Novagen and New England Biolabs, respectively. Ni-NTA agarose resin was supplied by QIAGEN. Amylose and *Strep*-Tactin Sepharose resins were from New England Biolabs and IBA, respectively. DNA modification enzymes and isopropyl β-D-thiogalactoside (IPTG) were from Promega. Oligonucleotides and molecular weight standards for native gel electrophoresis were supplied by Sigma. Bradford reagent was obtained from Bio-Rad.

### Construction of plasmids for expression of MutS mutants in *E. coli* and *P. aeruginosa*

MutSR272A and MutSR272E were obtained by changing the CGC (Arg) codon to GCC (Ala) and GAG (Glu) codons, respectively. MutSN279A was obtained by replacing the AAC (Asn) codon for the GCC (Ala) codon, and MutSF33A was obtained by changing the TTC (Phe) codon to the GCC (Ala) codon. These mutants were constructed by *Pfu* DNA polymerase amplification from pET15b-*mutS* (plasmid used for His_6_-tag expression in *E. coli*) (Pezza et al., 2002), pET15bStrep-*mutS* (plasmid used for *Strep* II-tag expression in *E. coli*) (Margara et al., 2016) or pEx-*mutS* (plasmid used for expression in *P. aeruginosa*) (Monti et al., 2012) using the following primers:

mutSPA-R272A-s

5’-CCTCGACGGCGCCAGCCGC<u>GC</u>CAACCT**C**GAGCTGGATATCAACCTCAGC-3’

mutSPA-R272A-as

5’-GCTGAGGTTGATATCCAGCTC**G**AGGTTG<u>GC</u>GCGGCTGGCGCCGTCGAGG -3’

mutSPA-R272E-s

5’-CCTCGACGGCGCCAG**T**CGC<u>GAG</u>AACCTGGAGCTGGATATCAACCTCAGC-3’

mutSPA-R272E-as

5’-GCTGAGGTTGATATCCAGCTCCAGGTT<u>CTC</u>GCG**A**CTGGCGCCGTCGAGG-3’

mutSPA-N279A-s

5’-GCCGCCGCAACCTGGAGCTGGA**C**ATC<u>GC</u>CCTCAGCGGTGGCCGCGAGAAC-3’

mutSPA-N279A-as

5’-GTTCTCGCGGCCACCGCTGAGG<u>GC</u>GAT**G**TCCAGCTCCAGGTTGCGGCGGC-3’

mutSF33A-s

5’-GATGTTCTATCGCATGGGCGA**T**<u>GC</u>**A**TACGAGCTGTTCTACGAGGAC-3’

mutSF33A-as

5’-GTCCTCGTAGAACAGCTCGTA**T**<u>GC</u>**A**TCGCCCATGCGATAGAACATC-3’

Changed nucleotides to introduce the point mutations are underlined. Silent changes to create *XhoI*, *Nru*I or *Nsi*I sites for the R272A, R272E and F33A mutations, respectively, and to destroy a *EcoR*V site for N279A are in bold. Following cycling, the product was treated with *Dpn*I to digest template DNA and mutated plasmids were transformed in *E. coli* DH5α.

### Expression and purification of recombinant proteins

Recombinant His_6_-tagged proteins were purified from *E. coli* BL21 (λDE3) using a Ni-NTA resin according to the manufacturer’s instructions. Maltose-binding protein (MBP) and *Strep* II-tagged proteins were expressed in *E. coli* DH5α and purified using Amylose and *Strep*-Tactin resins, respectively, as previously described (Margara et al., 2016, Miguel et al., 2008). All proteins were purified with a purity >95% as determined by SDS-PAGE. Protein concentration was determined by Bradford assay using BSA as a standard and aliquots were stored at -70 °C. Concentrations of tetrameric MutS proteins (MutS, MutSR272A, MutSR272E, MutSN279A and MutSF33A) and dimeric MutS mutants (MutSR842E and MutSΔ798) were expressed as tetramer and dimer molarity, respectively. β clamp and MutL were expressed as dimer molarity. MBP-HTH/Cter and MBP-Cter were expressed as tetramer molarity.

### DNA substrates preparation

DNA substrates were prepared by annealing the 37 bp oligonucleotide 5’-ATTTCCTTCAGCAGATAG<u>T</u>AACCATACTGATTCACAT-3’ with the following oligonucleotides: 5’-ATGTGAATCAGTATGGTT<u>A</u>CTATCTGCTGAAGGAAAT-3’, 5’-ATGTGAATCAGTATGGTT<u>G</u>CTATCTGCTGAAGGAAAT-3’, 5’-ATGTGAATCAGTATGGTT<u>A</u>CTATCTGC-3’ or 5’-ATGTGAATCAGTATGGTT<u>G</u>CTATCTGC-3’ to obtain dsDNA, GT-dsDNA, pDNA and GT-pDNA, respectively. The other DNA substrates used in this work are showed in Supplementary Data. Annealing reactions were carried out by heating to 100 °C for 5 min in 10 mM Tris-HCl (pH 7.4), 100 mM NaCl and then slow cooling at 25°C over 2 h.

### Native polyacrylamide gel electrophoresis

Native polyacrylamide gel electrophoresis assays were carried out by incubating affinity-purified proteins in the presence or absence of DNA oligonucleotides in binding buffer (20 mM Tris-HCl, pH 7.4, 10 mM MgCl_2_, 1 mM dithiothreitol, 100 mM NaCl, 150 µM ADP and 100 µg*/*ml BSA) for 20 min at 30°C. Samples were applied to native polyacrylamide gels prepared in Tris-borate buffer and run at 4°C. Gels were stained with ethidium bromide and/or coomassie blue. The intensity of protein or DNA bands was quantified using the Gel-Pro analyzer software (Media Cybernetics, Silver Spring, MD, USA). The relative mobility of protein-DNA complexes was compared to the standards: thyroglobulin (670 kDa), apoferritin (443 kDa), β-amylase (200 kDa), and bovine albumin (66 kDa). Migration of each standard band was plotted versus its molecular weight, and the slope of the resulting line, calculated by linear regression, was used to determine the approximate molecular weight of protein-DNA complexes.

### Electrophoretic mobility-shift assays

DNA binding reactions were performed by incubating various amounts of MutS proteins (15-4000 nM) with biotin labelled DNA oligonucleotides (10 nM) in binding buffer for 20 min at 30°C. Samples were applied to native polyacrylamide gels prepared in Tris-borate buffer and run at 4°C. Gels were fixed in 50 % (v/v) isopropanol and 5 % (v/v) acetic acid for 15 min, washed in water for 30 min and incubated in PBS containing 0.1% Tween-20 and 0.1 ng/ml IRDye 800CW Streptavidin for 1 h. After washing three times in PBS plus 0.1% Tween-20 for 10 min, gels were scanned in an Odysseey infrared imaging (LI-COR Bioscience) instrument and subjected to densitometry with the Odyssey software. Free DNA band intensities were quantified, and used to calculate the fraction of DNA bound and free MutS concentrations. Saturation binding curves were obtained by plotting the fraction of DNA bound versus free MutS concentrations. For the derivation of the apparent dissociation constant (K_D_), data points were fitted to a sigmoidal curve using the Sigma Plot software. The sigmoidal equation was Y= Y_0_ + Y_max_ (X^n^ / X^n^ + K_D_^n^), where Y is the fraction of DNA bound, Y_max_ is the maximum fraction of DNA bound, Y_0_ is the background signal, X is the concentration of free MutS, K_D_ is the apparent dissociation constant and n is the Hill-coefficient.

### Surface plasmon resonance

Surface plasmon resonance analysis was performed using a BIAcore T100 instrument (GE Healthcare) at 25°C. Biotinylated DNA substrates were immobilized on a streptavidin-derivatized sensor chip. The 37 bp template strand had a biotin at the 5’ end, and thus all DNA substrates were similarly oriented over the whole chip surface. Soluble proteins were diluted in binding buffer without BSA, and injected over chip surfaces at a flow rate of 30 µl min^−1^ for 180 s. Then, binding buffer was flown over for 220 s. The chip was regenerated with binding buffer containing 0.005% Tween-20 for pDNA, GT-dsDNA and dsDNA; and binding buffer amended with 4.5 M MgCl_2_ for GT-pDNA. SPR data were analyzed using Biacore T100 Evaluation software (Biacore, version 3.2). Dissociation constants (*K*_D_) for MutS-DNA interactions were determined by kinetic analysis fitting to a 1:1 Langmuir model, or by plotting the maximum response attained at each MutS concentration and fitting data to sigmoidal or hyperbolic curves. The equation for the sigmoidal curve was Y= Y_0_ + Y_max_ (X^n^ / X^n^ + K_D_^n^) and for the hyperbolic curve was Y= (Y_max_ X)/(K_D_ + X) + Y_0_ X, where Y is the maximum response, Y_max_ is the maximum signal, Y_0_ is the background signal, X is the concentration of MutS, n is the Hill-coefficient and K_D_ is the apparent dissociation constant.

### Circular dichroism

Far-UV CD spectra of 2 μM MutS proteins preincubated alone or with 4 µM DNA substrates in 6 mM Tris-HCl, pH 7.4, 7.5 mM NaCl and 0.28% glycerol were recorded using a Chirascan CD spectroscope (Applied Photophysics). Spectra were collected in triplicate from 250 nm to 190 nm at 25°C. The prediction of the proportion of the secondary structural element contents was performed using the BeStSel server available on the Web (http://bestsel.elte.hu/) (Micsonai et al., 2018).

### Limited proteolysis

2 μM MutS was preincubated alone or with 4 µM DNA substrates in binding buffer without BSA for 1 h at 4°C. Then, mixtures were reacted with trypsin (7.3-13 µg/ml) for 10 min at 37 °C. Proteolysis reactions were quenched by the addition of sample buffer and boiled for 10 min. The proteolytic products were separated on 8% SDS-polyacrylamide gel and coomassie blue stained. The full length-MutS band intensity was quantified using the Gel-Pro analyzer.

### ATP challenge assay

MutS-DNA complexes were assembled by incubating MutS (250 nM) with DNA oligonucleotides (50 nM) in binding buffer for 10 min at 30°C. Then, ATP (0.01-4000 nM) was added and reaction mixtures were incubated for 15 min at 30°C. Samples were applied to native polyacrylamide gels prepared in Tris-borate buffer and run at 4°C. Gels were stained with ethidium bromide and the intensity of free DNA bands was quantified using the Gel-Pro analyzer software. The effective concentration of ATP for 50% MutS dissociation (EC_50_) was determined by plotting the percentage of DNA bound against ATP concentrations and fitting the curve to the following equation: Y= Y_0_ + (Y_max_ EC_50_)/(X + EC_50_), where Y is the percentage of DNA bound, Y_max_ is the maximum percentage of DNA bound, Y_0_ is the background signal and X is the concentration of ATP.

### ATPase activity

To measure the ATPase activity of MutS and the mutant versions, these proteins (20-1280 nM) were individually mixed with 200 µM ATP in binding buffer. To determine the effect of DNA on the ATP hydrolysis catalyzed by MutS, reaction mixtures contained 40 nM MutS, 1.6 µM DNA and ATP (3-400 µM). Reactions were initiated by the addition of MutS and incubated for 20 min at 37°C. The liberation of phosphate from ATP was determined by using the malachite green assay. Briefly, the malachite green dye mixture (0.09% malachite green dye, 1.48% ammonium molybdate and 0.17% Tween 20) was added in a 1:4 ratio to reactions and incubated for 15 min at room temperature prior to being read at 612 nm. The standard curve was measured using NaH_2_PO4 solution.

### Docking and molecular dynamics simulations setup

MutS putative DNA binding sites were evaluated using the DNA binding predictor BindUp (http://bindup.technion.ac.il/) (Paz et al., 2016). For molecular docking, rigid 3D structural models of the DNA substrates were generated using Make-na (http://structure.usc.edu/make-na/) (Sim et al., 2012) and 3D-DART (van Dijk and Bonvin, 2009). Molecular docking between the MutS crystal structure (PDB: 1E3M) and DNA substrates was performed using HDOCK server (Yan et al., 2017). The docked structure of MutS and GT-pDNA, obtained by HDOCK server, was used as the starting structure of molecular dynamics (MD) simulations in solvent using GROMACS (Van Der Spoel et al., 2005). MD simulations were performed using the amber99sb force field (Hornak et al., 2006) and Tip3p water. An octahedral box was used, ensuring a solvent shell of at least 10 Å around the solute and resulting in a total of 109700 water molecules. The system was neutralized with 90 sodium ions that were placed at random positions. Simulations employed periodic boundary conditions and electrostatic interactions were treated using the particle-mesh Ewald algorithm (Essmann et al., 1995) with a real space cutoff of 10 Å. Lennard-Jones interactions were truncated at 10 Å. A Verlet cutoff-scheme was used. The system was initially subjected to 50.000 energy minimization steps using steepest descendent, followed by 50 ps of equilibration using position restraints in the protein backbone. Finally, 60 ns of production MD with no restraints were performed. Temperature (300 K) was maintained constant using a V-rescale temperature coupling, an isotropic pressure (1 bar) with a Berendsen coupling and a 2 fs time step. LINKS constraints were applied to all bonds involving hydrogens. Hydrogen bonds formed between MutS and DNA were analyzed using gmx hbond. MutS-DNA figure renders were obtained using VMD (Humphrey et al., 1996).

### Dot far-Western blotting

To perform dot far-Western blotting assays, affinity-purified proteins were spotted onto Protran nitrocellulose membranes (0.45 μm, GE Healthcare). After being dried, the membranes were blocked for 1 h at room temperature in blocking buffer (10 mM Tris-HCl, pH 7.4, 150 mM NaCl, 1 mM EDTA, 0.05% Triton X-100 and 5% milk), and then incubated with a His_6_- or *Strep* II-tagged protein binding partner in the same buffer overnight at 4°C. After washing, the membranes were incubated with a mouse anti-His_6_ monoclonal antibody (1/10,000, Sigma) or a mouse anti-Strep II monoclonal antibody (1/10,000, IBA) for 3 h at room temperature, washed, and then incubated for 1 h with IRDye 800CW-conjugated goat anti-mouse antibody (LI-COR Bioscience). Blots were scanned on an Odyssey infrared imager instrument (LI-COR Bioscience) and subjected to densitometry with the Odyssey software.

### Estimation of spontaneous mutation rates

Spontaneous mutations in different target genes on the *P. aeruginosa* chromosome were measured by estimating mutation rates to resistance to ciprofloxacin (*nfxB*, *gyrA* and *parC* are mutated), rifampicin (*rpoB* is mutated) and amikacin (*mexZ* is mainly mutated).

Mutation rates were determined by the modified Luria-Delbruck fluctuation test (Luria and Delbruck, 1943). Independent cultures (30) were obtained as follows: *mutS*-deficient PAO1 cells freshly transformed with the derivatives of pEx were inoculated in LB medium containing 20 μg/ml gentamicin in the presence of 10 µM IPTG (∼500 cells/ml) and grown to exponential phase. After cultures reached late-exponential phase (∼1 - 5 x 10^8^ cells/ml), aliquots from successive dilutions were plated onto LB agar containing 20 μg/ml gentamicin to determine the number of viable cells and onto LB containing 0.5 μg/ml ciprofloxacin, 50 μg/ml rifampicin or 10 μg/ml amikacin to select resistant cells. Colonies were scored after 24-48 h. The Ma-Sandri-Sankar (MSS) maximum-likelihood method was applied to estimate the number of mutants (m) using the MSS algorithm (Sarkar et al., 1992). Then, the mutation rate (μ) was calculated with the Luria–Delbruck equation μ=m/Nt, where Nt corresponds to the final number of cells in cultures (Luria and Delbruck, 1943). rSalvador program (Zheng, 2017) was used to calculate μ and 95% confidence limits.

### Western blot analysis

Total soluble protein extracts were performed by resuspending cells in 20 mM Tris-HCl, pH 7.4, 0.5 M NaCl, 15% glycerol amended with 0.2 mg/ml lysozyme, 1 mM phenylmethylsulfonyl fluoride and 1 mM benzamidine, and incubating for 1 h on ice. After four sonication (2 min) and freeze/unfreeze cycles, intact cells were removed by centrifugation at 9000 *g* for 20 min and the extracts were stored at -20 °C. Proteins (50 μg) were separated by SDS-PAGE (8 %). The resolved proteins were transferred to nitrocellulose membranes (0.22 μm, Sigma) and probed with a mouse anti-Strep II monoclonal antibody (1/10,000, IBA) for 3 h at room temperature, washed, and then incubated for 1 h with IRDye 800CW-conjugated goat anti-mouse antibody (LI-COR Bioscience). Membranes were scanned on the Odyssey infrared imager instrument (LI-COR Bioscience) and subjected to densitometry with the Odyssey software.

## ACKNOWLEDGEMENTS

This work was supported by grants from Consejo Nacional de Investigaciones Científicas y Técnicas [PIP 20130100360]; Secretaría de Ciencia y Tecnología de la Universidad Nacional de Córdoba [33620180101008CB]; and Agencia Nacional de Promoción Científica y Técnica [PICT 2017-1857]. Funding for open access charge: Agencia Nacional de Promoción Científica y Técnica [PICT 2017-1857].

## AUTHOR CONTRIBUTIONS

Investigation, M.I.I.B., L.M.M., S.D.C., M.M.F., G.G.M. and V.M.; Formal analysis, E.L.M., V.M. and C.E.A.; Funding acquisition, C.E.A. and M.R.M.; Writing – Original Draft, M.R.M.; Writing – Review & Editing, M.R.M.; Conceptualization, M.R.M.; Methodology, M.M.F., V. M. and M.R.M.

## SUPPLEMENTARY TABLES

**Table S1.** Docking scores for the interaction between *E. coli* MutSΔ800 and DNA structures^1^.

| Structure number | 1 | 2 | 3 | 4 | 5 | 6 | 7 | 8 | 9 | 10 |
| --- | --- | --- | --- | --- | --- | --- | --- | --- | --- | --- |
| **GT-dsDNA** | -201.4 | **-180.9** | **-171.7** | -166.9 | **-164.0** | -162.3 | -159.9 | **-158.5** | -158.3 | -157.4 |
| **dsDNA** | -195.6 | **-194.1** | **-191.6** | -185.3 | -182.3 | **-174.5** | -174.0 | **-165.3** | **-164.4** | **-163.9** |
| **GT-pDNA** | **-256.2** | -245.8 | **-213.3** | -198.9 | -196.8 | **-195.1** | **-194.3** | **-192.9** | -192.0 | -190.9 |
| **pDNA** | **-243.3** | **-218.6** | -217.5 | **-215.1** | -207.3 | -202.4 | -199.1 | -196.9 | **-192.0** | **-190.9** |
| **ssDNA** | -293.2 | -290.6 | -283.9 | -283.1 | -281.7 | -79.52 | -277.1 | -273.9 | -272.6 | -270.3 |

<sup>1</sup>Docking scores values for the top ten scored structures of the *E. coli* MutSΔ800-DNAs complexes are shown. DNA structures docked in the DNA binding site I or II are highlighted in red or blue, respectively.

**Table S2.**
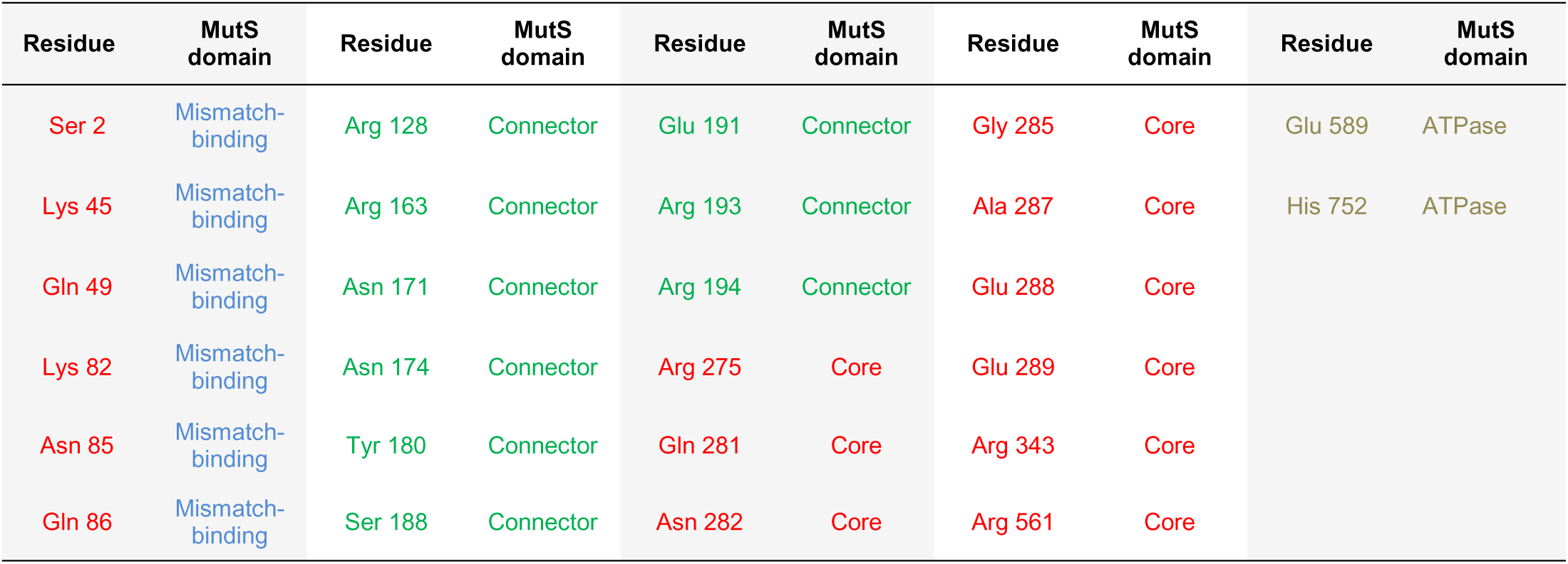
Residues involved in hydrogen bonds.

## SUPPLEMENTARY FIGURES

**Figure S1.**
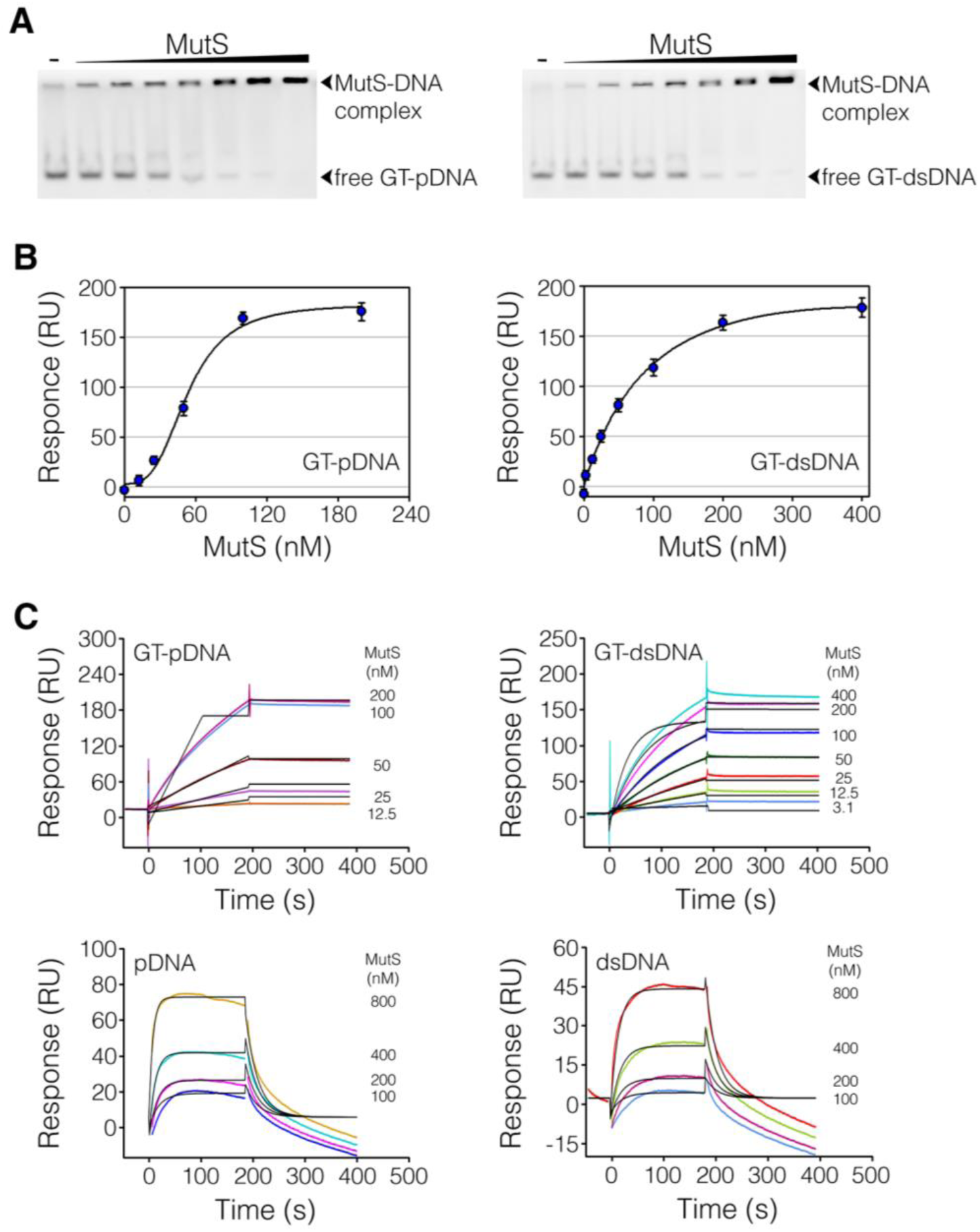
MutS interaction with replication DNA substrates. (A) Analysis of MutS interaction with DNA substrates by EMSA. MutS (5, 15, 30, 60, 125, 250 and 500 nM) was incubated with a fixed concentration of biotinylated DNA oligonucleotides (10 nM), and reaction products were analyzed on native polyacrylamide gels. Free and bound DNAs were detected by staining gels with infrared dye-labeled streptavidin. Representative images of gels are shown for MutS binding to GT-pDNA and GT-dsDNA. (B and C) SPR measurements of MutS interaction with biotinylated DNA immobilized on a streptavidin-derivatized sensor chip. (B) Determination of the K_D_ for MutS binding to GT-pDNA and GT-dsDNA using maximum responses for different protein concentrations. (C) Kinetic fitting of MutS binding to DNA substrates to a 1:1 Langmuir model. Different protein concentrations are represented by different colors and fitted kinetics are shown in black lines. Sensorgrams of MutS display solvent correction. Representative reference-subtracted curves are shown (n=3).

**Figure S2.**
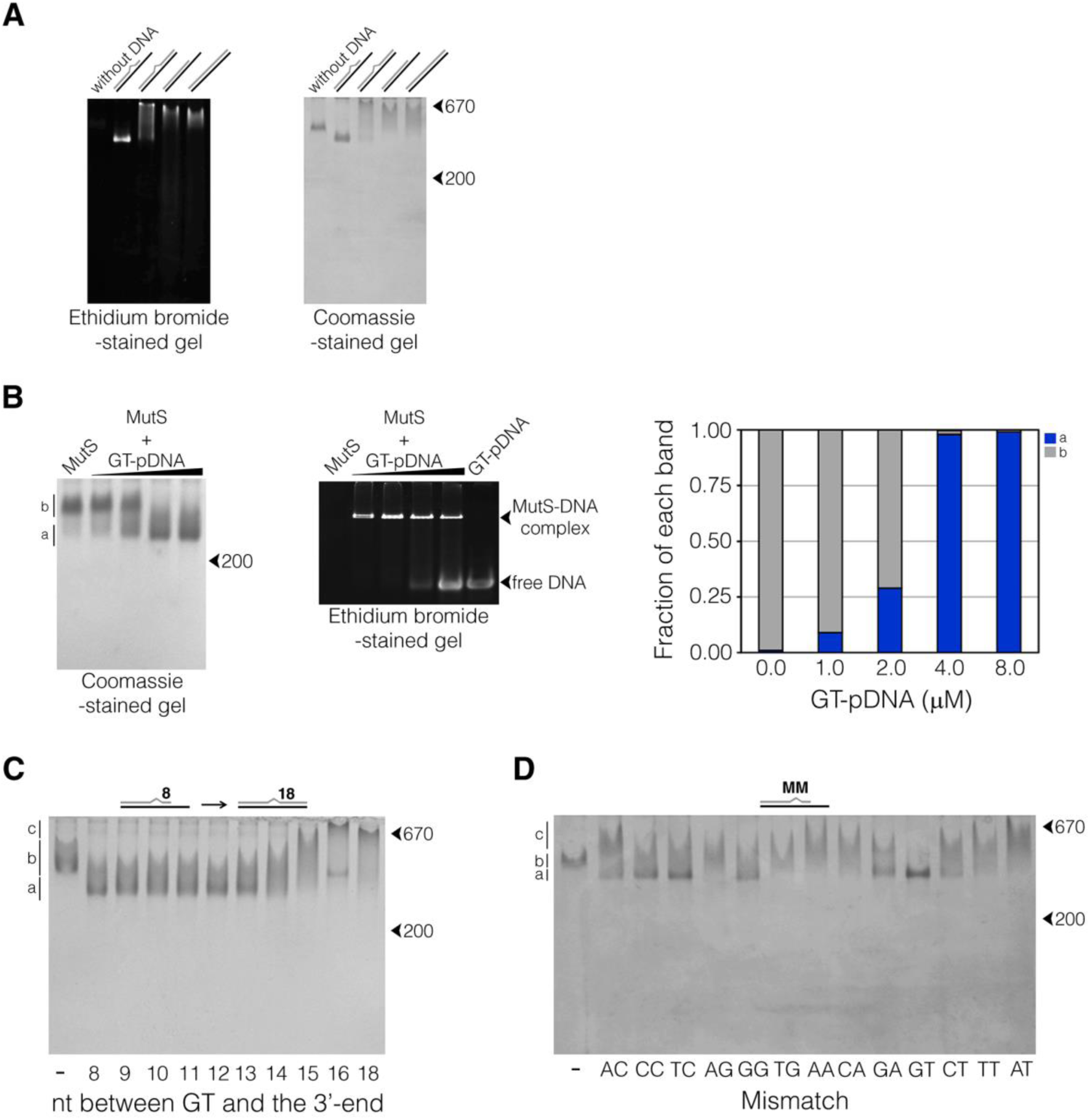
Electrophoretic migration of DNA-bound MutS under native gels. (A) MutS (2 μM) was individually incubated alone or with DNA substrates (4 μM). Reaction mixtures were analyzed on native polyacrylamide gels. Gels were stained with ethidium bromide (left panel) and subsequently with coomassie blue (rigth panel) to detect DNA and protein, respectively. Bands that showed both ethidium bromide and coomassie blue staining were identified as the MutS-DNA complexes. (B) MutS (2 μM) was incubated alone or with increasing GT-pDNA concentrations (1, 2, 4 and 8 μM), and reaction mixtures were analyzed on native polyacrylamide gels. Representative images of ethidium bromide- (10%) and coomassie blue-stained (6.5%) gels are shown. The intensity of a and b bands was quantified in each line to calculate the relative fraction of each band. Data represented the means of at least triplicates. Standard deviations were <5% of the mean. (C) Representative image of a coomassie blue-stained gel showing the mobility of MutS (2 μM) bound to GT-pDNA structures (4 μM) containing 8 nt to 16 nt between GT and the 3’-OH terminus of the primer strand. Control reactions without DNA (-) and GT-dsDNA (18 nt from the GT) were included. (D) Representative image of a coomassie blue-stained gel showing the mobility of MutS (2 μM) bound to pDNA structures (4 μM) containing the twelve different mismatches (MM) located 8 nt from the 3’-OH terminus of the primer strand. Control reactions without DNA (-) and pDNA (AT) were included. Migration of molecular weight markers, thyroglobulin (670 kDa) and β-amylase (200 kDa), is shown with arrows.

**Figure S3.**
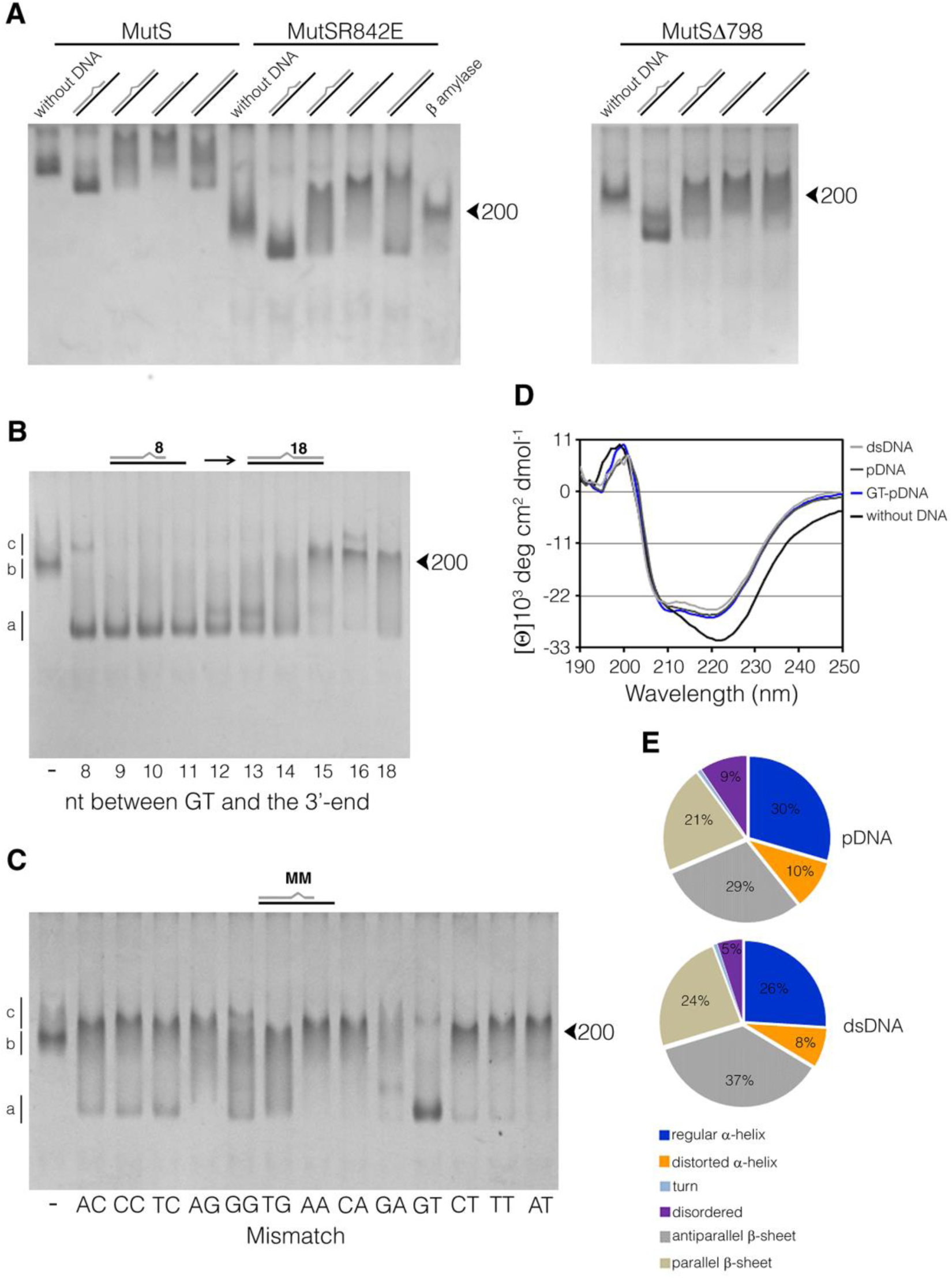
Rearrangement of the MutS structure upon binding to mismatched replication DNA substrates. (A) Representative image of a coomassie blue-stained gel showing the mobility shift of DNA-bound MutS, MutSR842E and MutSΔ798 relative to the free proteins. MutS (2 μM), MutSR842E (4 μM) and MutSΔ798 (4 μM) were individually mixed with DNA substrates (4 μM), and reaction products were analyzed on native polyacrylamide gels. (B) Representative image of a coomassie blue-stained gel showing the mobility of MutSR842E bound to GT-pDNA structures containing 8 nt to 16 nt between GT and the 3’-OH terminus of the primer strand. Control reactions without DNA (-) and GT-dsDNA (18 nt from the GT) were included. (C) Representative image of a coomassie blue-stained gel showing the mobility of MutSR842E bound to pDNA structures containing the twelve different mismatches (MM) located 8 nt from the 3’-OH terminus of the primer strand. Control reactions without DNA (-) and pDNA (AT) were included. Migration of β-amylase (200 kDa) is shown with an arrow. (D) Far-UV CD spectra of 2 μM MutS in the absence (black line) or presence of 4 μM GT-pDNA (blue line), pDNA (dark-grey line) and dsDNA (light-grey line). The plotted spectra are the average of three independent scans. Contributions of buffer and DNA to ellipticity were subtracted. (E) Fraction of secondary structures estimated from CD spectra using the BeStSel web tool.

**Figure S4.**
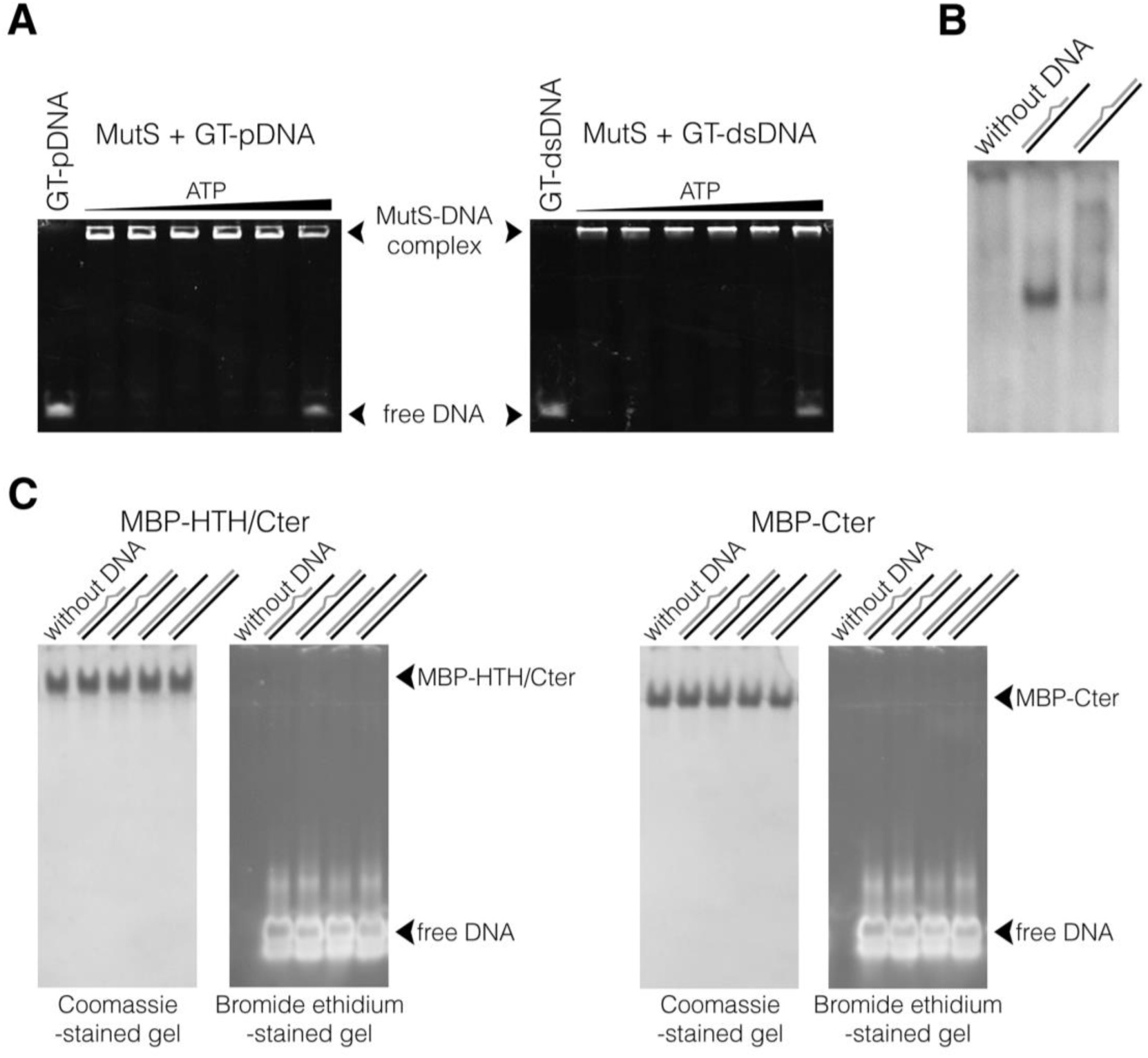
(A) MutS association with mismatched DNA structures upon ATP challenge. MutS (250 nM) bound to DNA oligonucleotides (50 nM) was incubated with increasing concentrations of ATP, and reaction products were analyzed on native polyacrylamide gels. We tested ATP concentrations in a range of 0.01-4000 µM. MutS dissociated from DNA at ATP concentrations higher than 100 µM. Representative images of ethidium bromide-stained gels showing the release of MutS prebound to GT-pDNA and GT-dsDNA substrates when challenged with 0.01, 0.1, 1, 10, 100 and 1000 µM ATP . (B) MutS from *E. coli* (2 μM) was individually incubated alone or with DNA substrates (4 μM) in 20 mM Tris-HCl pH 7.4, 10 mM KCl, 5 mM MgCl_2_, 1 mM DTT and 50 µg/ml BSA. Reaction mixtures were analyzed on native polyacrylamide gels (run in TAE 1 x buffer). Representative image of a coomassie blue-stained gel showing that GT-pDNA induced a higher mobility of *E. coli* MutS relative to the free protein, which did not migrate into the gel. *E. coli* MutS bound to GT-dsDNA showed a diffuse migration, with a small fraction (∼18%) of the complex migrating near the band corresponding to *E. coli* MutS-GT-pDNA. (C) The C-terminal domain (Cter, residues 799-855) alone or along with the contiguous helix-turn-helix domain (HTH, residues 764-855) were fused to the maltose binding protein to generate MBP-Cter and MBP-HTH/Cter, respectively. Both fusions (13.5 µM) were incubated with DNA substrates (16 µM), and reaction products were analyzed on native polyacrylamide gels. No binding was observed between fusions and DNA substrates since DNA-protein complexes were not detected under these conditions. This result could indicate that the HTH and Cter domains do not interact with DNA; however, further experiments are needed to establish this fact. Representative ethidium bromide-(right panels) and coomassie blue-stained (left panels) gels are shown.

**Figure S5.**
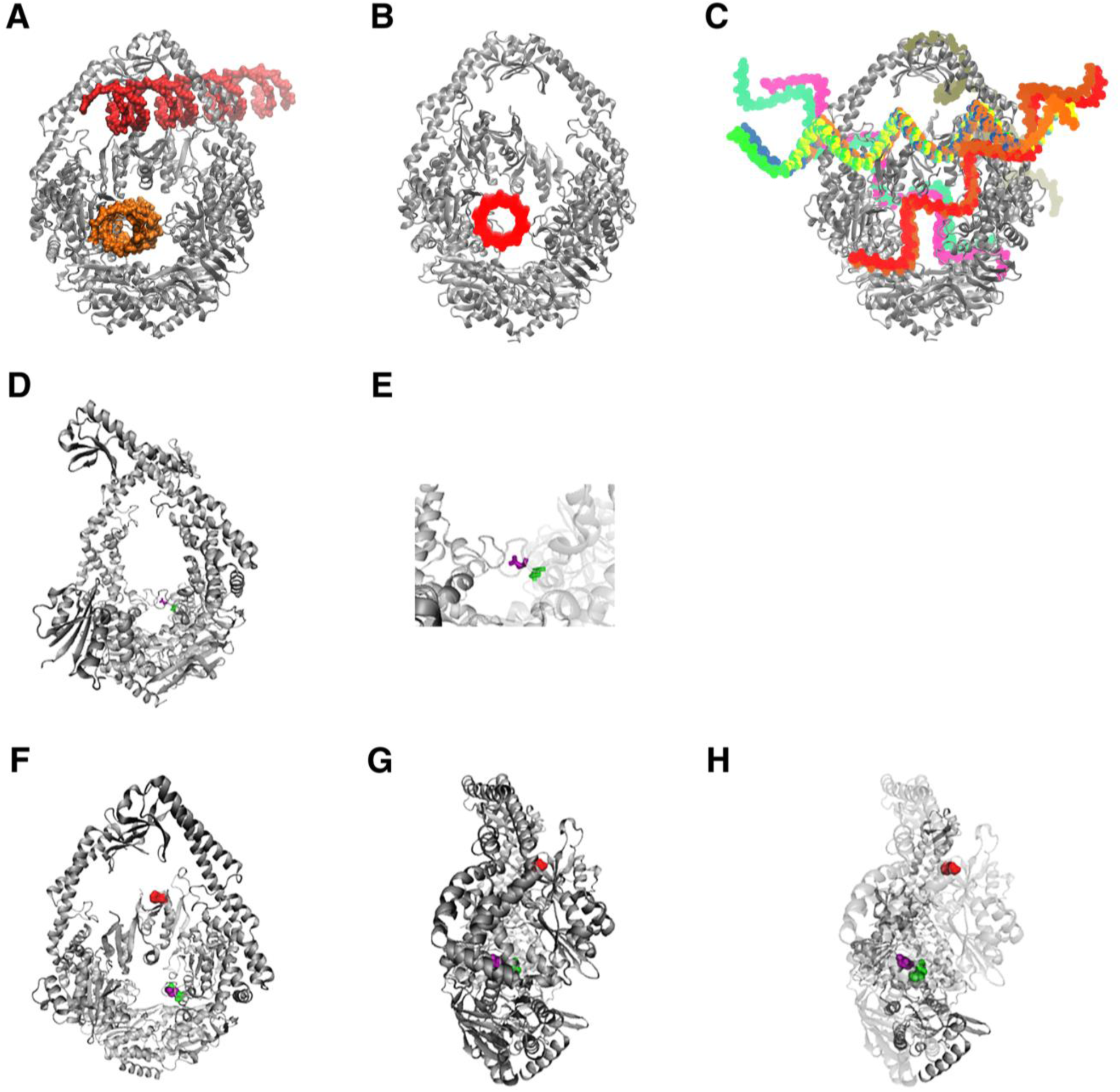
MutS contains a novel putative DNA interaction site for DNA replication structures. (A-C) Docking simulations of the *E. coli* MutSΔ800 (PDB entry: 1E3M) interaction with dsDNA, pDNA and single-stranded DNA. MutSΔ800 is represented by ribbons diagrams (front view, grey). DNAs are shown in a space-filling model. (A) Structures of MutSΔ800 with docked dsDNA showed that this DNA can interact with the DNA binding sites I and II. (B) pDNA docked specifically in the DNA binding site II, whereas the single-stranded DNA did not contact the upper and lower channels of MutSΔ800 (C). (D and E) Front view of *E. coli* MutS Δ800 sliding clamp (PDB entry: 5AKD; ribbon representation; gray) showing Asn 282 and Arg 275 (stick representation; magenta and green, respectively). (F-H) Front and lateral view of *E. coli* MutSD835R (PDB entry: 3ZLJ; ribbon representation; gray) showing the DNA binding residues Phe 36, Asn 282 and Arg 275 (surface representation; red, magenta and green, respectively).

**Figure S6.**
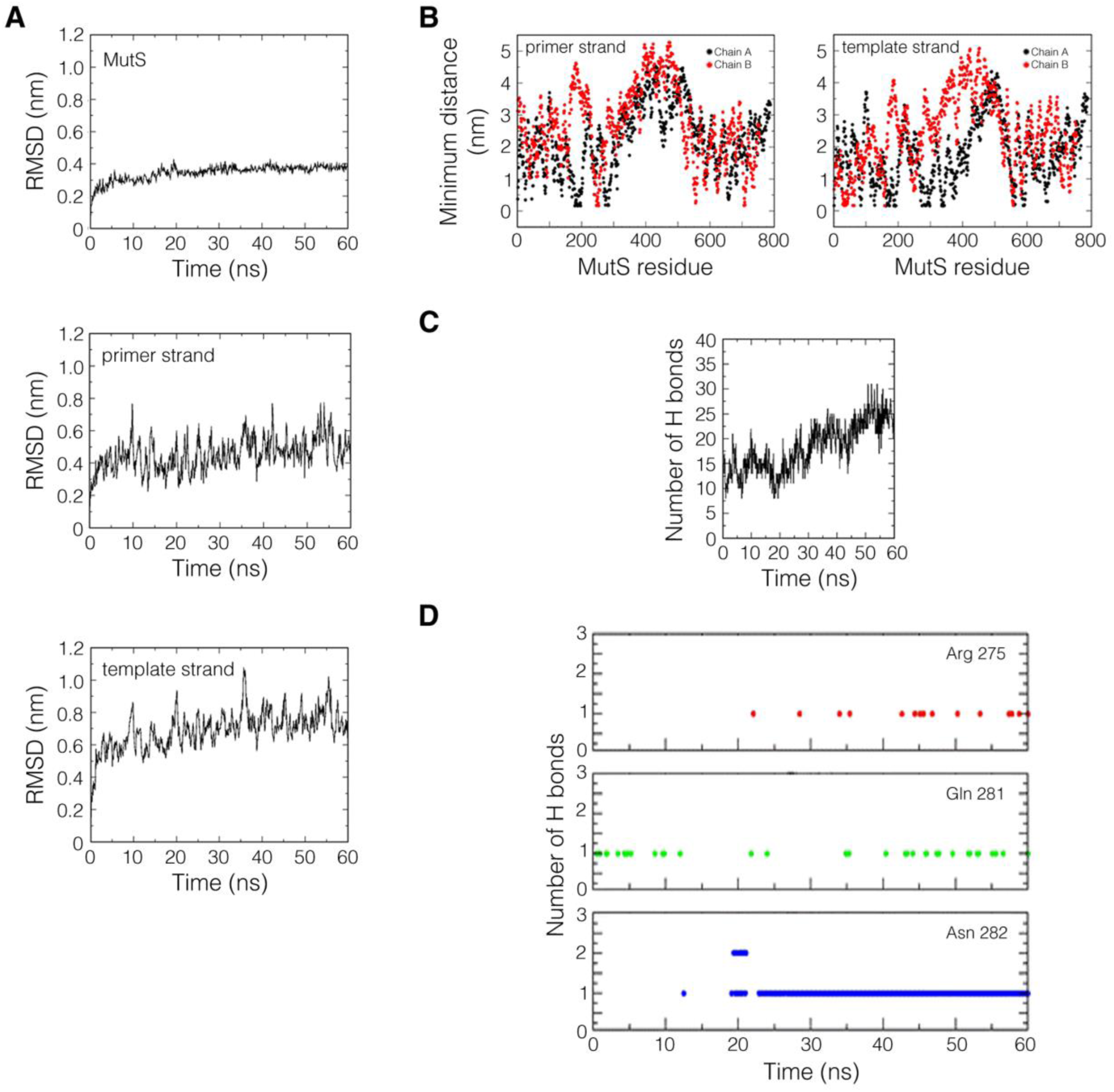
Molecular dynamics simulations of the docked MutS-GT-pDNA complex. (A) Root mean square deviations (RMSD) of MutS and GT-pDNA backbone chains (primer and template strands). (B) Minimum distances between MutS chain A (black dots) and chain B (red dots) residues and the primer and template strands of GT-pDNA. (C) Number of hydrogen bonds formed between MutS and GT-pDNA during the course of simulation. (D) Number of hydrogen bonds formed between Arg 275, Gln 281 and Asn 282 residues and GT-pDNA.

**Figure S7.**
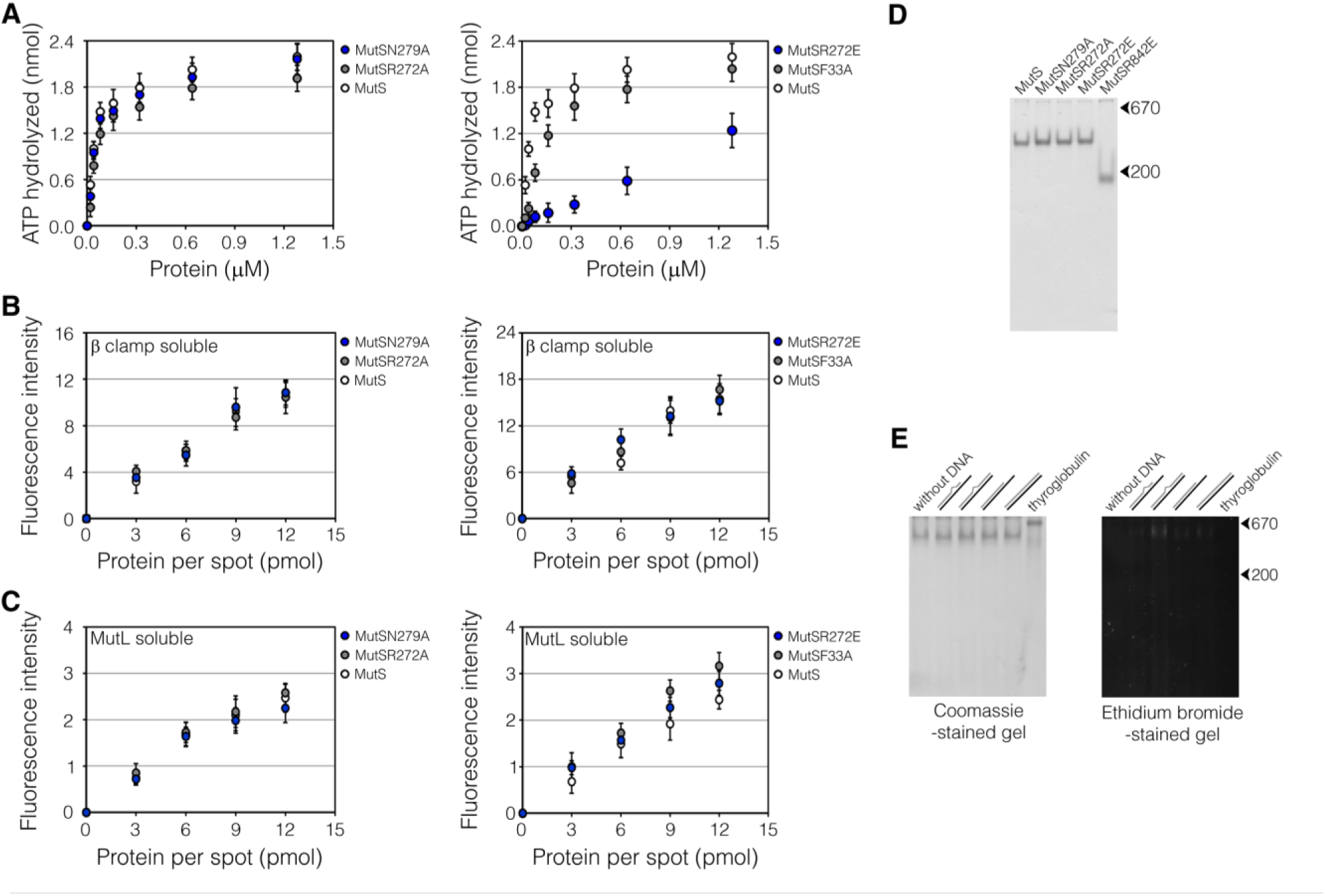
Effect of N279A, R272A and R272E substitutions on MutS function. (A) ATP hydrolysis by wild-type and MutS mutants was measured by quantifying the free phosphate released in parallel reactions containing MutS proteins (0.020-1.28 µM) and ATP (200 µM). (B and C) Spots of *Strep* II-tagged MutS, MutSN279A, MutSR272A, MutSR272E and MutSF33A (3, 6, 9 and 12 pmol) on nitrocellulose membranes were incubated with soluble 0.25 μM His_6_-tagged MutL or β clamp. Binding of soluble proteins was detected using a mouse anti-His_6_ monoclonal antibody. The fluorescence intensity of the spots was quantified using the Gel-Pro analyzer software. Data represent mean ± standard deviation (n=3). (D) Representative image of a coomassie blue-stained gel showing the mobility of the MutSN279A, MutSR272A and MutSR272E mutants compared to the migration of tetrameric MutS, and dimeric MutSR842E. (E) MutSF33A (2 μM) was individually incubated alone or with DNA substrates (4 μM). Reaction mixtures were analyzed on native polyacrylamide gels. Representative images of ethidium bromide- and coomassie blue-stained (6.5%) gels are shown. Migration of molecular weight markers, thyroglobulin (670 kDa) and β-amylase (200 kDa), is shown with arrows.

**Figure S8.**
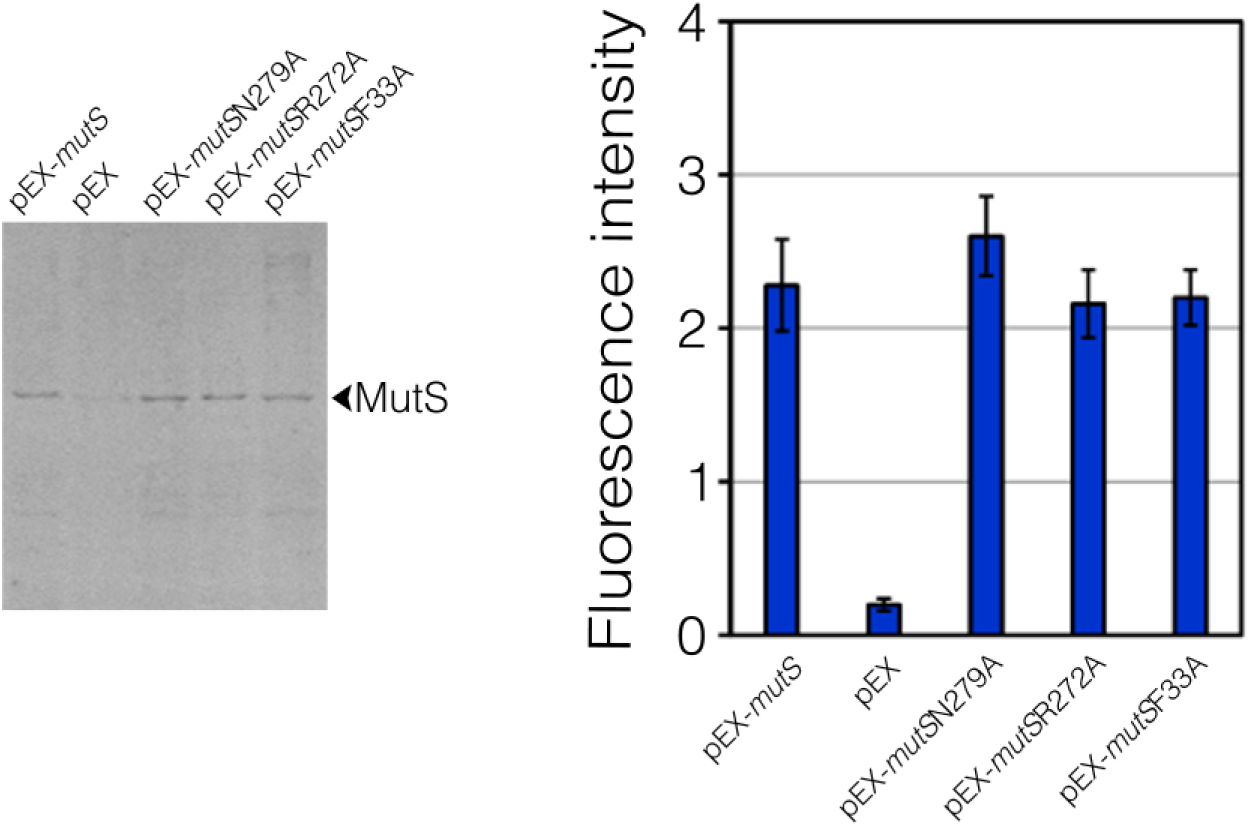
Detection of ectopically expressed MutS proteins by Western blotting. MutS and mutant versions were expressed as Strep II-tagged fusions using the IPTG-inducible pEx plasmid. Expression levels of these proteins were examined in PAO1 *mutS* cells carrying empty pEx, pEx-*mutS*, pEX-*mutS*N279A, pEx-*mutS*R272A and pEx-*mutS*F33A after induction with 10 µM IPTG by Western blotting using an anti-Strep II antibody. A representative Western blot membrane is shown (right-panel). The fluorescence signal of protein bands was measured, and data represent the means ± standard deviations (n=3).

## BindUp predicted DNA-binding sites

### *E. coli* MutS domains (Lamers et al., 2000)

Mismatch-binding domain: residues 2–115

Connector domain: residues 116–266

Core domain: residues 267–443 and 504–567

Clamp domain: residues 444–503

ATPase domain: residues 568–765

Helix–turn–helix (HTH) domain: residues 766–800

C-ter domain: 801-853

### BindUp predicted positive surfaces in full length *E. coli* MutSD835R bound to a GT-dsDNA structure (PDB ID: 3ZLJ)

Largest Positive Patches in Monomer A:

<u>Patch 1</u>: PRO615 ASN616 MET617 GLY618 GLY619 ILE695 HIS728 TYR729 PHE730 LEU747 PHE758 MET759 HIS760 LYS769 SER770 TYR771 GLY772 LEU773 ALA774 VAL775 ALA776 LEU778 VAL781 PRO782 LYS783 GLU784 VAL785 ILE786 LYS787 ARG788 ARG790 GLN791 LYS792 LEU793 ARG794 GLU795 SER823 PRO824 LEU837 PRO839 ARG840 GLN841 ALA842 GLU844

<u>Patch 2</u>: GLN129 ASP130 ASN131 LEU132 THR147 ARG154 ARG156 LEU157 GLN171 ARG172 THR173 ASN174 GLU261 ARG274 **<u>ARG275</u>** THR280 GLN281 ALA292 MET306 LYS308 ARG309 TRP310 HIS312 MET313 VAL315 VAL320

<u>Patch 3</u>: LEU18 LYS21 ALA22 PRO25 GLU26 TYR41 ASP42 LYS45 ARG46 GLN49 LEU55 LYS57 PRO65 PRO67

Largest Positive Patches in Monomer B:

<u>Patch 1</u>: SER770 TYR771 LEU773 ALA774 ILE786 LYS787 ARG790 GLN791 LEU793 ARG794 SER798 SER823 PRO824 VAL826 ASP833 PRO834 ARG835 LEU837 GLN841 LEU852

<u>Patch 2</u>: ASP461 THR462 LYS464 VAL465 GLN476 ILE477 SER478 GLY480 GLN481 LEU495 LYS496 ASN497 ALA498

<u>Patch 3</u>: LEU132 ARG156 ARG172 GLU261 ARG309 HIS312 MET313

### *Thermus aquaticus* MutS domains (Obmolova et al., 2000)

Mismatch-binding domain: residues 1–118

Connector domain: residues 132–245

Core domain: residues 247–385 and 514–540

Clamp domain: residues 406–513

ATPase domain: residues 543–765

### BindUp predicted positive surfaces in *Thermus aquaticus* MutSΔ768 bound to an unpaired T-dsDNA structure (PDB ID: 1EWQ)

Largest Positive Patches in Monomer A:

<u>Patch 1</u>: ALA259 HIS558 VAL561 PHE567 ASN585 ALA587 GLY588 SER590 THR591 PHE592 ARG594 GLN595 ILE626 GLY627 ASP662 GLU663 ALA694 HIS726

<u>Patch 2</u>: TYR449 LEU450 TYR457 ARG465 PRO466 GLN474 TYR476 THR477 LEU478 PRO479 GLU480 LYS482 GLU483 ARG486

<u>Patch 3</u>: ALA131 TYR133 GLU154 PHE171 ARG172 HIS173 ARG174 TYR244 PHE249 PHE263 GLU264 THR271 SER274 ARG286 LEU287 GLN289 HIS294

Largest Positive Patches in Monomer B:

<u>Patch 1</u>: PRO14 PRO15 LEU16 LEU17 GLN19 VAL36 GLY37 ASP38 THR59 HIS60 LYS61 THR62 SER63 LYS64 PHE66 GLN97 VAL98 VAL108 ARG110

<u>Patch 2</u>: LEU260 LEU266 HIS558 VAL561 PHE567 ASN585 GLY588 LYS589 SER590 THR591 PHE592 ARG594 GLN595 ILE626 GLY627 ALA628 ALA694 HIS726

<u>Patch 3</u>: GLU154 GLY213 LEU217 ARG221 PHE241 ARG242 PHE243 TYR244 PHE249 ARG286 SER290 HIS294

### Human MutSα (MSH2-MSH6) domains (Warren et al., 2007)

Mismatch-binding domain: MH2 residues 1–124; MSH6 residues 362-518

Connector domain: MSH2 residues 125–297; MSH6 residues 519-717

Core domain: MSH2 residues 300–456 and 554–619; MSH6 residues 718-934 and 1009-1075

Clamp domain: MSH2 residues 457–553; MSH6 residues 935-1008

ATPase domain: MSH2 residues 620-855; and MSH6 residues 1076-1355

### BindUp predicted positive surfaces in human MutSα (MSH2-MSH6Δ341) bound to a GT-dsDNA structure (PDB ID: 2O8B)

Largest Positive Patches in MSH2:

<u>Patch 1</u>: LYS90 LEU94 GLY126 LEU128 SER129 GLN130 GLU132 ILE134 LEU135 PHE136 GLY137 ASN138 ASN139 ILE145 GLY146 VAL166 SER168 ILE169 ARG171 ILE194 PRO196 LYS197 LEU277 GLU278 LEU279 PHE286 PHE289 THR335 PRO336 GLN339 ARG340 ASN343 GLN344 LYS347 PHE384 PRO385 ASN388 ARG389 LEU390 ALA391 LYS392 LYS393 GLN395 ARG396 GLN402 ASP403 TYR405 ARG406 PHE468 VAL589 GLN593

<u>Patch 2</u>: ARG219 GLY220 GLY221 ILE304 ARG308 ALA309 SER323 HIS639 CYS641 MET672 GLY674 LYS675 SER676 THR677 TYR678 ARG680 GLN681 VAL712 GLY713 ASP748 GLU749 ARG752 VAL802 TYR815

<u>Patch 3</u>: MET26 PRO27 PRO30 THR32 THR33 ARG35 ARG99 VAL100 GLU101 TYR103 LYS122 ALA123 SER124 THR225 ARG227 LYS228 ALA230 ALA272 LYS275 PHE276 GLU278

Largest Positive Patches in MSH6:

<u>Patch 1</u>: LEU740 ASN741 PHE746 HIS1096 THR1133 GLY1134 PRO1135 ASN1136 MET1137 GLY1139 LYS1140 SER1141 ARG1145 ARG1176 LEU1177 GLY1178 THR1189 GLU1193 GLU1196 ASP1213 GLU1214 ARG1217 GLY1218 THR1247 HIS1248 TYR1249 HIS1250 SER1251 VAL1253 MET1267 ALA1268 CYS1269 MET1270 THR1284 ALA1293 CYS1294 LYS1296 SER1297 TYR1298 GLY1299 PHE1300 ASN1301 ALA1302 VAL1312 LYS1315 GLY1316 HIS1317 ARG1318 LYS1319 ALA1320

<u>Patch 2</u>: TYR366 ASN404 SER405 CYS406 THR407 PRO408 GLY409 MET410 ARG411 LYS412 GLN415 ILE416 VAL429 GLY430 LYS431 PHE432 TYR436 HIS437 TRP456 GLY460 PHE461 PRO462 ILE464 ARG468 TYR469 GLN485 GLU487 PRO489 MET492 GLU493 CYS496 MET499 HIS501 ILE502 SER503 LYS504 TYR505 ARG507 VAL508 VAL509 ARG510 ARG511 ILE513 TYR845 GLU846

<u>Patch 3:</u> TYR524 SER525 VAL526 LEU527 GLU528 LYS824 1 VAL828 PRO831 LYS833 SER834 HIS837 ASP839 SER840 TYR850 SER851 LYS853 LYS854 ASP857 PHE933 SER935 ASP938 LYS1030 MET1033 ARG1034 ARG1035 TYR1038

## DNA SUBSTRATES USED IN THE PRESENTE WORK

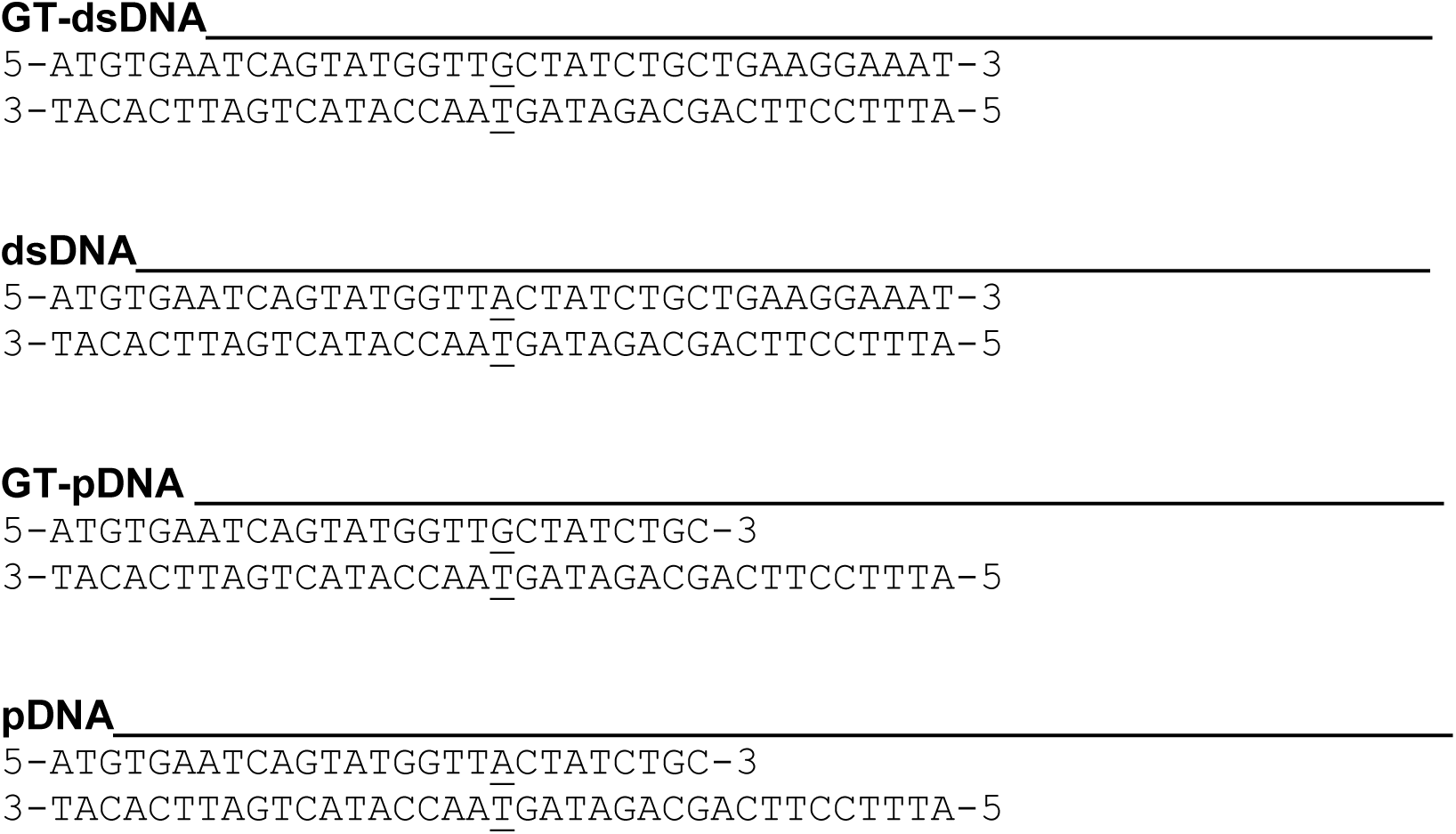

### DNA substrates used to test the effect of extending the 3’-OH end of the primer strand in the GT-pDNA

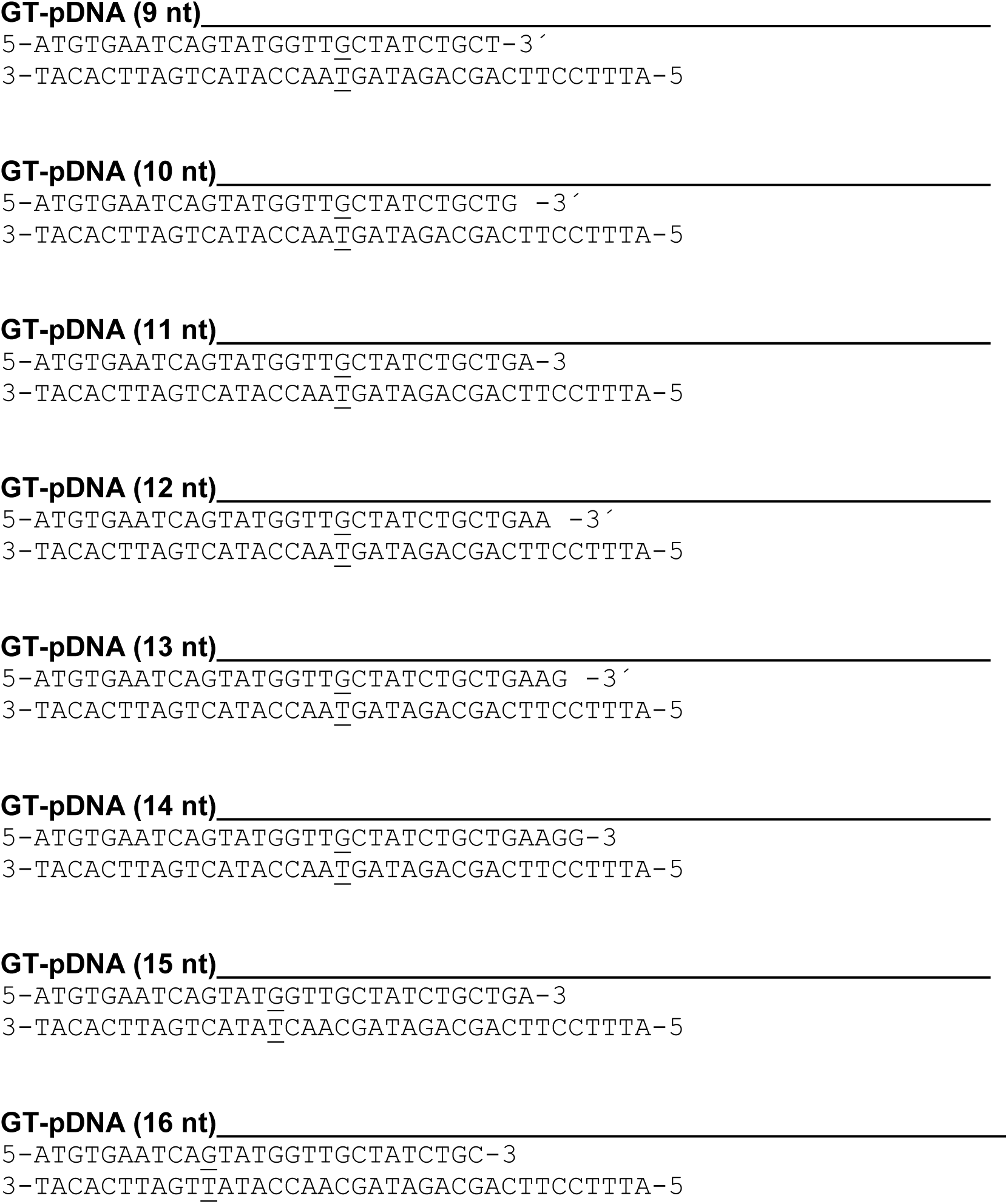

### DNA substrates used to test the effect of different mismatches in the pDNA

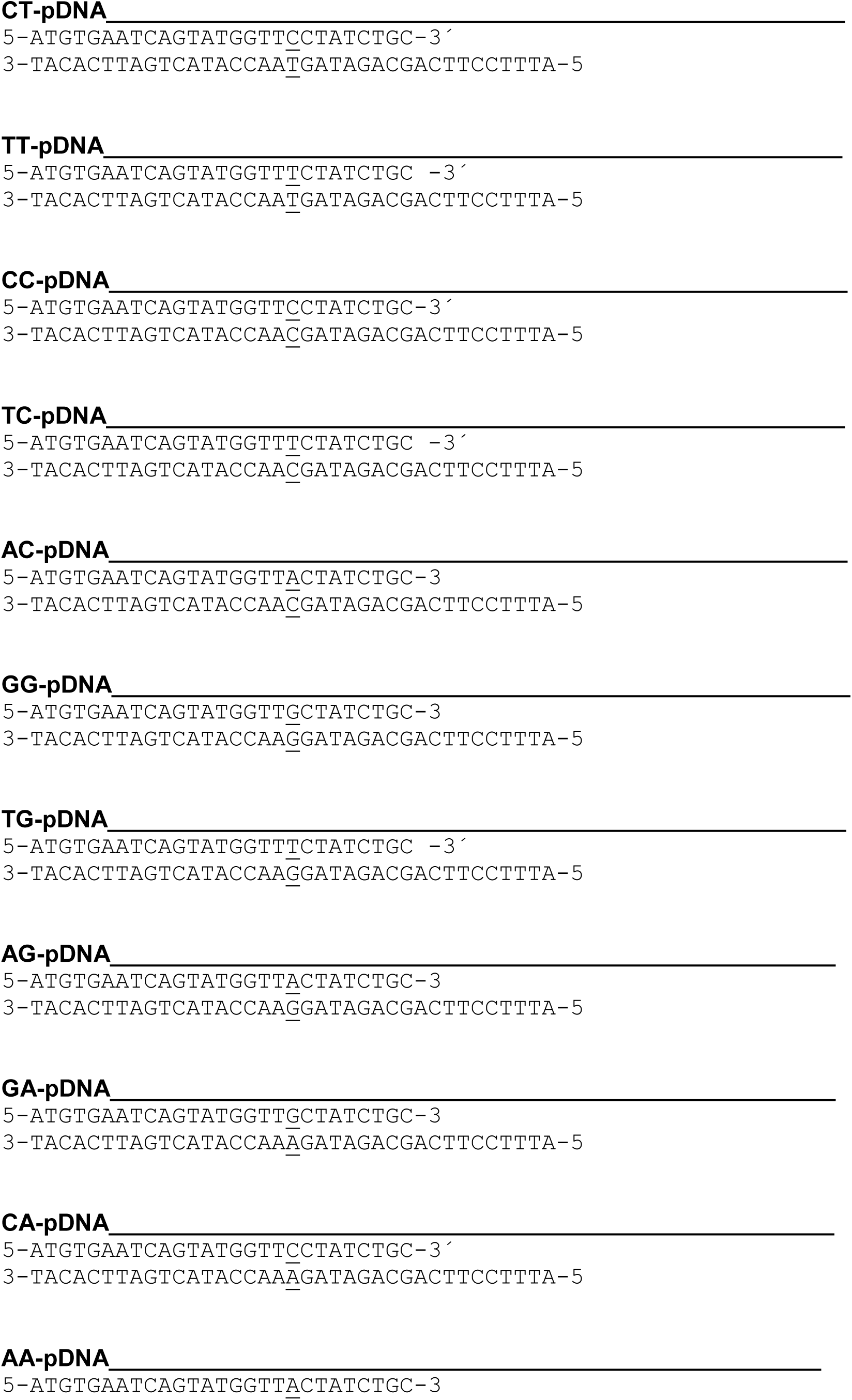
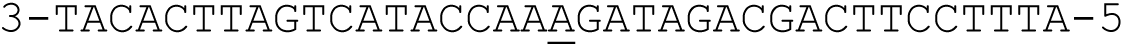

